# Evaluating Variant Effect Prediction Across Viruses

**DOI:** 10.1101/2025.08.04.668549

**Authors:** Sarah Gurev, Noor Youssef, Navami Jain, Aarushi Mehrotra, Sarrah Rose Mikhail Leung, Abigail Jackson, Debora Marks

**Affiliations:** Systems Biology, Harvard Medical School; FutureHouse; Broad Institute; Bioinformatics and Integrative Genomics, Harvard Medical School; Harvard-MIT Health Sciences and Technology

## Abstract

Viruses are a major threat to global health due to their rapid evolution, extensive diversity, and frequent cross-species transmission. Although advances in machine learning and the expanding availability of sequence and structural data have accelerated large-scale mutation effect prediction, viral proteins, and particularly fast-evolving antigenic proteins, pose unique biological and data-related challenges that may limit model performance. We introduce EVEREST, a curated dataset for evaluating model performance on (i) forecasting real-world viral evolution (31 clades across 4 viruses) and (ii) concordance with lab-based deep mutational scanning assays (45 proteins, *>*340,000 variants). Using EVEREST, we show that state-of-the-art protein language models trained across the protein universe substantially under-perform on viral proteins relative to alignment-based models trained on homologous proteins. This under-performance persists even in low-sequence regimes, as is the case during a novel viral outbreak. We develop calibrated reliability metrics to quantify confidence in model predictions where no evaluation datasets exist. For more than half of the WHO-prioritized pandemic-threat viruses, current models fail to produce reliable predictions, highlighting the urgent need for more data or new modeling approaches. Together, these findings reveal key factors driving model under-performance and provide actionable recommendations for improving viral mutation effect prediction in preparation for current and future outbreaks.

## Introduction

Viral pathogens pose a persistent and growing threat to global health. Their rapid evolution and high mutation rates often undermine the durability of vaccines and therapeutics, while increasing human–animal interactions heighten the risk of zoonotic spillovers. In response to these concerns, the World Health Organization (WHO) recently identified 40 priority and prototype RNA viruses across 16 families with high pandemic potential and limited medical countermeasures ^1^. Accurately forecasting how these viruses evolve could transform surveillance and therapeutic design by identifying immune or treatment-evading strains early, from the first time they are sequenced. In the case of SARS-CoV-2, computational approaches were able to flag Variants of Concern months before their designation by the WHO ^2,3^. While both experimental and computational tools exist to predict the functional impact of mutations, experimentally testing large numbers of variants remains a significant challenge and laboratory settings capture specific phenotypes which often do not fully recapitulate real-world selective pressures. Despite these limitations, experimental assays have proven valuable for predicting the evolution of key SARS-CoV-2 mutations ahead of their emergence during the COVID-19 pandemic ^4,5^.

Computational approaches offer a scalable alternative that learn fitness constraints directly from evolutionary sequences. Some models estimate mutation effects from large-scale sequencing of a specific viral population, through clade-growth, mutation-counting, and other phylogenetic approaches ^6–11^. While valuable, these methods are primarily retroactive, quantifying the fitness impact of mutations after they have emerged during viral evolution. The challenge for many viruses, including the WHO priority pathogens, is the ability to forecast the fitness effects of as-yet-unseen mutations, and before extensive surveillance sequencing is available. Other computational approaches, commonly used in protein design ^12–16^, overcome this challenge by learning fitness constraints from broader evolutionary sequences. These approaches differ by their input data, training objective, model scale and performance on different tasks ^17^. For example, alignment-based variant effect predictors (VEPs) learn from multiple sequence alignments (MSAs) of homologous proteins ^18–20^; Protein language models (PLMs) learn from large sets of sequences from across the tree of life ^21–26^; Inverse folding models are trained to predict an amino acid sequence given a three dimensional structure ^27,28^; and tree-based approaches look for conservation across subtrees of a protein family ^29^. Here, we focus on the most commonly used approaches for modeling viral evolution: alignment-based VEPs, PLMs, and hybrid models (Fig. 1).

**Figure 1:**
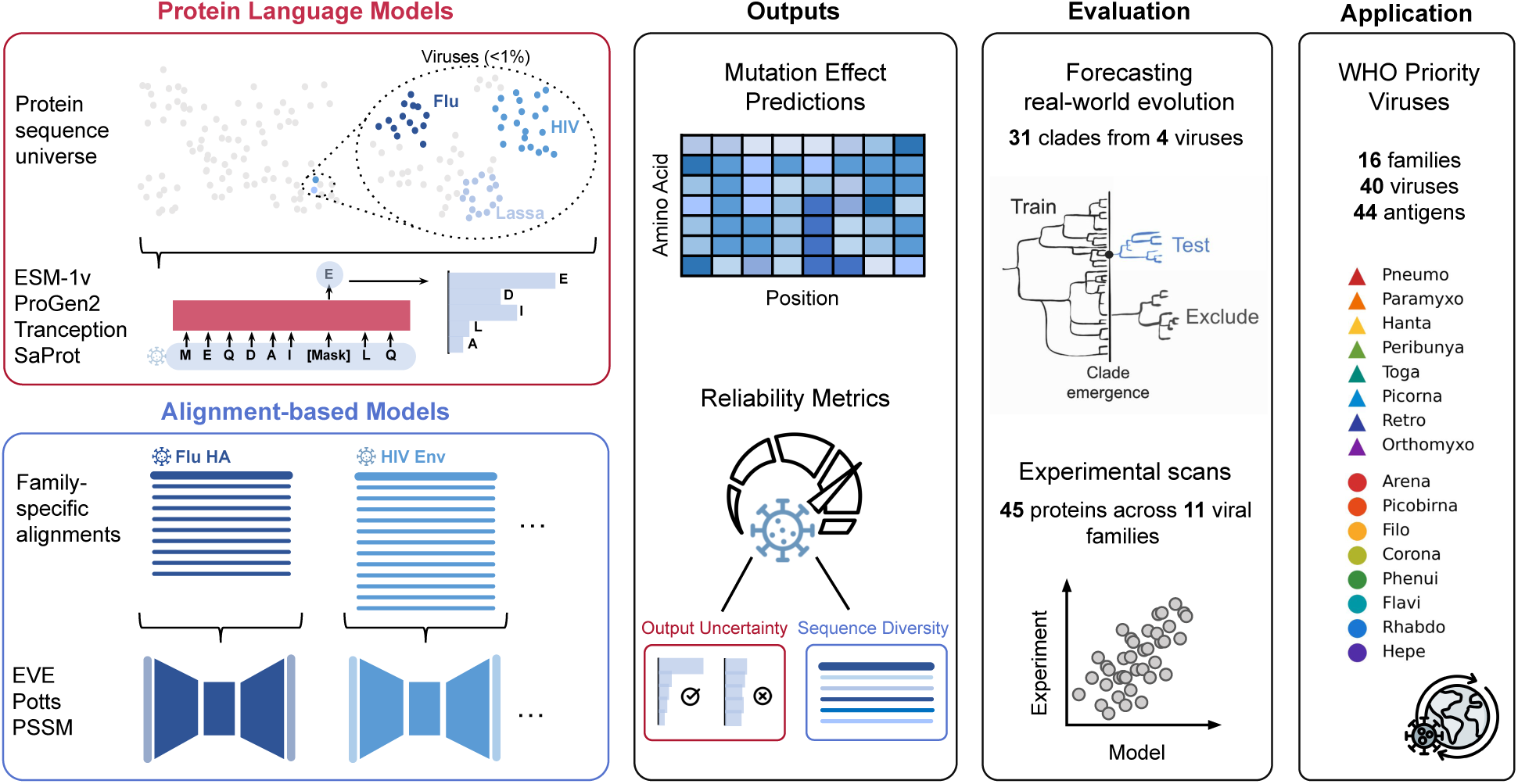
Comprehensive evaluation of variant effect prediction models for viral evolution. We evaluate the performance of four Protein Language Models (PLMs: ESM1v, ProGen2, Tranception, and SaProt), three alignment-based models (PSSM, Potts model, and EVE) and four hybrid models (VESPA, Tranception with MSA, TranceptEVE, and SaProt-EVE) using EVEREST, a curated dataset for viral tasks. This assessment includes large-scale experimental deep mutational scans (45 DMS across 11 viral families) and real-world viral evolution forecasting (31 clades across 4 viruses; SARS-CoV-2, HIV, and influenza H3N2 and H1N1). We calibrate per-protein reliability metrics to estimate model uncertainty in the absence of experimental data and apply this to quantify model reliability across 40 WHO-prioritized viruses.

Alignment-based models have shown success in modeling viral evolution, immune escape, and vaccine effectiveness, especially for highly-sequenced viruses like HIV and SARS-CoV-2^2,3,18–20,30–36^. Given a query protein, related proteins are retrieved from sequence databases and are aligned to form a MSA. This set of sequences is assumed to include positive examples enriched for the same properties as the query. Therefore, a density model of these homologous, “evolutionary” sequences can be used as proxy for fitness constraints. The generalizability of alignment-based VEPs to high-risk viral families is currently unknown and may be limited, as many of the high-risk viruses currently lack diverse sequence data. Moreover, these VEPs cannot accommodate insertions and deletions—common features of viral evolution ^37^—due to their reliance on fixed-length sequence representations.

PLMs offer an alignment-free alternative for modeling protein mutation effects. Trained on large corpora of sequences, these models can handle sequences of arbitrary length and, in theory, generalize across protein families ^21–26^. PLMs span a broad design space, ranging from single-sequence models to architectures that incorporate MSA information during inference or training ^22,38–40^, as well as hybrid or structure-aware models ^41–43^. Single-sequence PLMs, the most common class, are trained on millions of natural proteins across diverse families to learn masked amino-acid prediction tasks. Although PLMs now achieve state-of-the-art performance on many general protein benchmarks ^17^, their effectiveness on viral proteins has yet to be systematically evaluated, despite their increasing use in this domain ^2,44–46^.

Unique features of viral evolution challenge extrapolation of best modeling practices from evaluations on non-viral tasks ^17^. As obligate intracellular parasites, viruses evolve under strong diversifying selection, particularly in antigenic regions. As such, viral proteins (especially those in RNA viruses) mutate at rates far exceeding those of cellular organisms and exhibit high structural and functional plasticity ^47,48^ (Fig. S1A). Consequently, generalized substitution matrices such as BLOSUM often fail to capture the patterns of mutations in viral proteins ^2,49^. These unique viral features raise a key question: can existing models *reliably* forecast viral mutation effects and evolutionary trajectories—especially during the early stages of an outbreak when experimental data is limited or not yet available?

To address this gap, we developed EVEREST (Evaluating Viral variant Effect prediction with Reliability ESTimation), a comprehensive benchmark tailored to viruses. EVEREST includes evaluation on two key tasks: forecasting real-world viral evolution and predicting variant effects from viral deep mutational scans (DMS). Real-world forecasting provides the strongest test of model performance but evaluation on this task is only possible for a small number of extensively sequenced viruses, such as SARS-CoV-2, HIV, and seasonal influenza (H1N1 and H3N2). To enable broader evaluation across diverse viruses, we include evaluations across 45 DMS datasets spanning 11 viral families. Using EVEREST, we evaluate 11 state-of-the-art models, including alignment-based VEPs, PLMs and hybrid models, and introduce calibrated metrics that quantify model reliability.

Across the 40 WHO-prioritized viruses, we find that current state-of-the-art models frequently fail to generate reliable predictions for surface glycoproteins—the primary antigenic determinants (hereafter, “antigens”) critical to vaccine design and immune resistance—substantially limiting their immediate utility for pandemic preparedness. This systematic assessment reveals the underlying reasons existing models underperform on viral proteins and provides actionable guidelines for improving future architectures and training strategies. Together, these insights lay the foundation for developing models capable of early and accurate forecasting of viral evolution, a critical component of global pandemic preparedness.

## Results and Discussion

### EVEREST experimental and real-world forecasting benchmarks

The EVEREST benchmark includes two evaluation datasets, measuring two complementary tasks: forecasting real-world viral evolution (forecasting benchmark) and predicting variant effects measured by viral deep mutational scanning assays (DMS benchmark; Fig. 1). For the forecasting benchmark, we identified 31 clades across four well-characterized viral protein antigens—SARS-CoV-2 spike, HIV envelope, and influenza H3N2 and H1N1 hemagglutinins—and assessed models on their ability to predict emerging mutations and prioritize highly frequent variants during evolution (Fig. S1). For alignment-based models, we implemented chronological train–test splits, restricting training data to sequences available prior to each clade’s emergence. In contrast, we note that since PLMs cannot be retrained at this scale, there is unavoidable data leakage (where mutations seen in the test set are included in training), resulting in inflated performance in all but the most recent clades. To contextualize forecasting performance, we compare models to the predictive ability of high-throughput fitness DMS measurements ^5,50–53^, while acknowledging that not all DMS assays are intended to serve as forecasting tools.

For the DMS benchmark, we follow well-established practices for evaluating the ability of general protein design models to predict mutation effects and design functional proteins ^17,18^. We curated a viral DMS benchmark that includes 45 viral proteins from 11 viral families, and more than 340,000 mutations—vastly broadening viral diversity compared to our forecasting benchmark ^4,50,54–78^. Measured experimental phenotypes include general fitness properties (e.g., infectivity, replication, and cell entry) and specific traits (e.g., stability and expression; Fig. S1, Table S1). We exclude antibody escape assays due to the limited number of antibodies or sera tested for most viruses, which can result in apparent false positives where a model correctly predicts escape but the DMS assay does not capture it.

Using EVEREST, we evaluate three alignment-based methods (PSSM ^18^, Potts model ^18^, and EVE ^19^); three PLMs (ESM-1v ^21^, Tranception ^22^, and ProGen2^24^); one structure-aware PLM (SaProt ^41^); and four hybrid approaches (TranceptEVE ^38^, Tranception with MSA retrieval ^22^, VESPA ^79^, and SaProt-EVE). These models were chosen for their diverse architectures, training datasets, and high performance on prior general benchmarks (Table S2).

### Protein Language Models underperform on viral proteins

Protein language models (PLMs), deep learning models trained on large-scale protein sequence datasets, have achieved state-of-the-art performance on mutation effect prediction in general protein benchmarks ^17^. As a result, prior studies have assumed that PLMs would represent a paradigm shift over alignment-based models for viral evolution prediction ^44^. However, despite their success in other domains, we find that single-sequence PLMs underperform on viral proteins compared to proteins from other taxa, with most PLMs having less accurate predictions than even the simplest site-independent alignment-based model (Fig. 2A, dotted lines).

**Figure 2:**
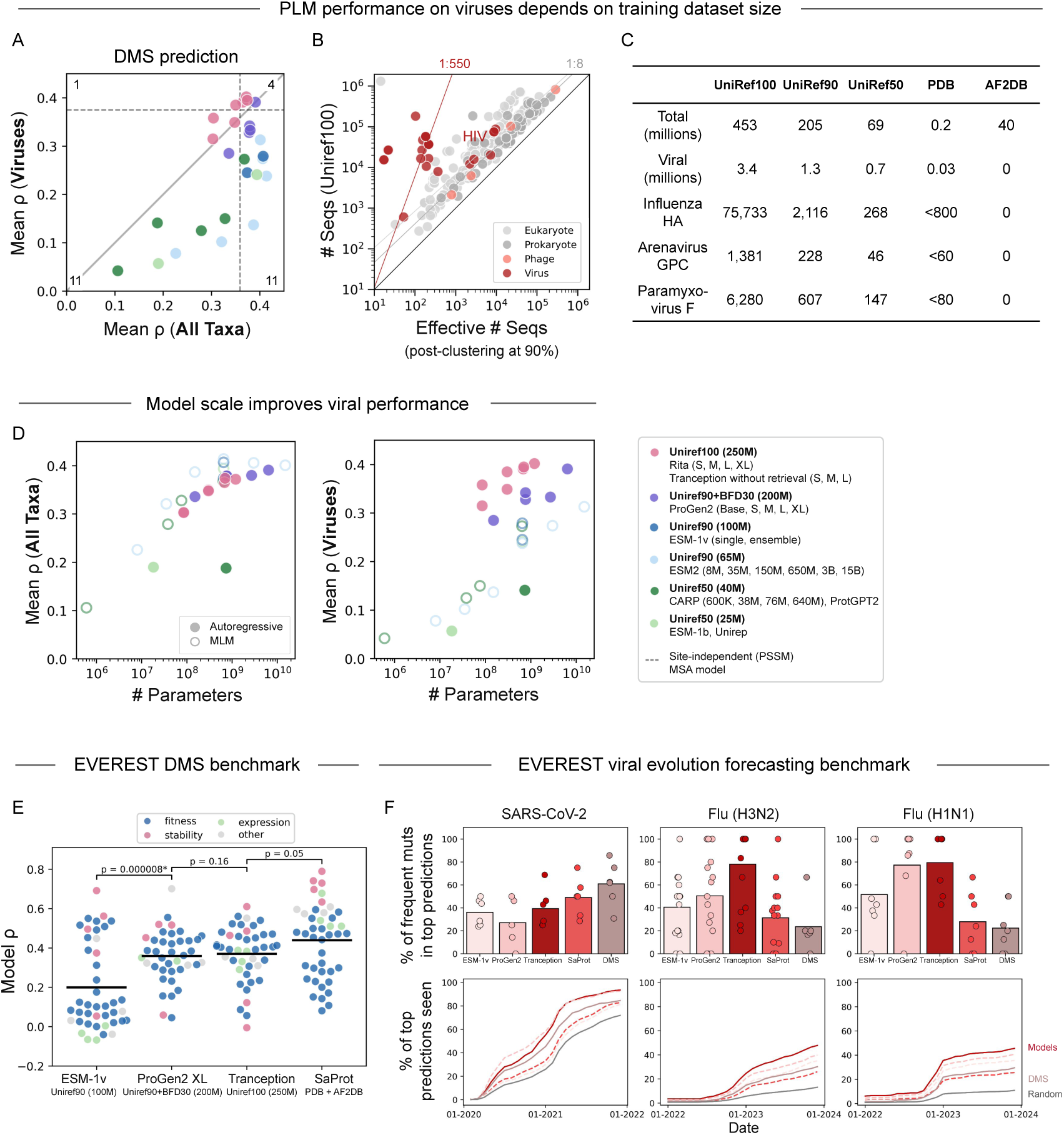
Evaluation of protein language models for viral evolution and mutation-effect prediction. **A.** Protein language models (PLMs) show reduced performance on viral proteins compared to other taxa. Performance is measured as the average Spearman correlation with experimental deep mutational scanning (DMS). Results include 218 DMS assays across all taxa versus *<* 20 assays from eukaryote-infecting viral proteins from prior benchmark ^17^. Dotted line indicates the average performance of alignment-based PSSM model, without any of the viral-specific alignment optimization discussed in the Alignment-based models section. See Table S3 for details. **B.** Sequence clustering disproportionately reduces viral diversity. While non-viral proteins maintain a ratio of 1 effective sequence (clustered at 90% identity) for every 8 raw (unclustered) sequences, most viruses exhibit a drastic reduction to 1:550. **C.** Viruses are underrepresented in PLM training, making up *<*1% of sequence databases. **D.** Unlike performance on other taxa (left) which saturates with model scale, viral performance (right) increases with both model size and training dataset volume. Each point denotes an individual PLM, colored by training dataset and objective (autoregressive vs. masked language modeling). **E.** PLM performance on EVEREST DMS benchmark, measured as Spearman correlation between model predictions and experimental DMS scores. Other PLMs show significant improvement over ESM-1v (*P <* 0.01; paired t-test). **F.** PLM performance on EVEREST viral forecasting benchmark. Top: percentage of highly frequent mutations captured by model predictions across 6 SARS-CoV-2 clades, 9 H1N1 clades and 15 H3N2 clades. Bottom: percentage of model-predicted mutations subsequently observed during evolution. Time-course analyses shown for representative clades—SARS-CoV-2 Wuhan (wild-type vaccine strain) and current flu vaccine strains for H1N1 (6B.1A.5a.2a.1), and H3N2 (3C.2a1b.2a2a.3a.1). Model predictions correspond to top 250 ranked mutations (see Fig. S7,S8 for alternate thresholds). Frequent mutations seen *>*1k times for flu and*>*10k times for SARS-CoV-2.

A likely driver of PLM underperformance on viral proteins is the relatively poor representation of viruses in PLM training datasets (*<*1% of training; Fig. 2B-C). PLMs are typically trained on clustered sequence databases, such as UniRef90 or UniRef50, which retain only one representative per sequence cluster with ≥90% or ≥50% identity, respectively. While clustering reduces redundancy and improves performance for well-sequenced protein families ^21^, it disproportionately filters out already rare viral sequences. While non-viral proteins maintain a ratio of 1 effective sequence (clustered at 90% identity) for every 8 raw (unclustered) sequences, most viruses exhibit a drastic reduction to 1:550 (Fig. 2B). This disproportionate downsampling is likely due to over-sequencing of narrow parts of the viral sequence landscape and the orders of magnitude higher mutation rates of viruses ^47^, leading to related sequences which more quickly evolve past the limits of homology detection (Fig. S1A). There are two notable exceptions to this downsampling trend: HIV and phages (Fig. 2B). HIV, with its extreme genetic variability and large sequencing efforts ^80^, has sufficient sequence diversity to allow for structural contact recovery from coevolutionary analysis, a property rarely observed in other viral antigens (Fig. S10, S5). Phages, as mostly double-stranded DNA viruses, have lower mutation rates than most eukaryote-infecting RNA viruses (Fig. S1A), resulting in higher number of identifiable homologs.

Two modeling choices tend to mitigate the poor representation of viral proteins in training datasets: (i) using larger models that may extract more informative features from sparsely represented proteins, and (ii) training on more comprehensive datasets with greater viral diversity. We find that increasing model size consistently improves accuracy on viral mutation effect prediction, in contrast to the performance plateau observed for non-viral proteins (Fig. 2D, Fig. S2) ^23,25,81,82^. Further supporting this, we find that ESM-2 models trained on the same dataset but spanning 8 million to 15 billion parameters show steady performance gains on viral proteins, as well as on non-viral proteins that are also poorly represented in training, with little to no improvement for well-represented proteins (Fig. S3).

Training dataset choice also affects performance in viral compared to non-viral proteins. PLMs trained on more comprehensive datasets (e.g., unclustered UniRef100) outperform those trained on clustered datasets (e.g., UniRef90, UniRef50) only for eukaryote-infecting viruses (Fig. 2A, S2, Table S3). For other taxa—including phages, which are not disproportionately downsampled in clustered databases (Fig. 2B)—models trained on UniRef90 or UniRef50 achieve superior performance compared with UniRef100-trained models (Fig. S2). The inclusion of metagenomic datasets (e.g., BFD30, BFD90, or OMG) provides minimal benefit for viral mutation effect prediction (Fig. S4). For example, ProGen2 trained on UniRef90+BFD30 performs similarly to the same model trained on UniRef90+BFD90 (Fig. S5B, S4), and both underperform compared to models trained on UniRef100, even when controlling for model size (Fig. 2D). Likewise, metagenomic sequences do not improve performance for SaProt-OMG relative to SaProt, despite the former being a substantially larger model (Fig. S5C, S4). Together, these results suggest that larger, unclustered datasets yield superior PLM performance on viral proteins and that metagenomic sequences do not compensate for the loss of natural evolutionary diversity introduced by clustering.

A confounding factor when interpreting training dataset-size effects is that models trained on the largest datasets (e.g., Tranception and ProGen2) employ an autoregressive objective, predicting each residue conditioned on all previous residues, rather than masked language modeling (MLM), which predicts only a small fraction of randomly masked residues (Fig. 2D, Table S3). It remains unclear whether performance gains arise primarily from increased dataset size or from the more challenging autoregressive objective, which introduces longer contiguous spans of uncertainty and may promote better generalization. Disentangling these contributions represents an important direction for future investigation.

We also examined other features that may contribute to performance. While most PLMs use input windows of 1,024 tokens, the effective context length is typically closer to half the training length ^83,84^, which is smaller than the length of most viral antigens. Nonetheless, context length alone is unlikely to explain the observed performance disparities. ProGen2 Base and ProGen2 Medium share identical architectures and training data but differ only in context length (2,048 vs. 1,024 tokens); the longer-context model performs slightly worse on viral proteins (Spearman’s *ρ* = 0.328 vs. 0.342; Table S3) ^24^. We also find that ensembling across multiple random seeds provides modest improvements across taxa, including viruses (Table S3).

Taken together, these observations suggest that larger PLMs trained on larger, unclustered datasets (e.g,. Tranception-L and ProGen2-XL) are better suited for viral applications in zero-shot settings than PLMs trained on smaller datasets with an MLM objective (e.g., ESM-2^44–46^).

#### Tranception outperforms other PLMs on EVEREST

Using our EVEREST viral forecasting and DMS benchmarks, we further evaluate four PLMs trained on different databases and with different training objectives: ESM-1v (ensemble) trained on UniRef90 (100M sequences), ProGen2-XL trained on UniRef90 + BFD30 (200M), Tranception-L trained on UniRef100 (250M) and the structure-aware PLM SaProt trained on the PDB + AF2DB (40M structures). All models share transformer-based architectures: both SaProt and ESM-1v use a BERT-style MLM objective, whereas Tranception and ProGen2 are autoregressive models ^21,22,24,41^.

ProGen2-XL, Tranception, and SaProt had comparable performance across the 45 DMS assays, significantly outperforming ESM-1v (Fig. 2E, *P* < 0.0001; paired t-test). ESM-1v only outperformed ProGen2 and Tranception on proteins with high numbers of homologous sequences in Uniref90 (i.e., HIV and phage proteins; Fig. 2B, S5A,F). SaProt’s strong performance on DMS assays is primarily driven by experiments measuring stability, expression, and other properties where structural context may be especially informative (Fig. 2E, Fig. S5A). The comparable performance of ProGen2-XL and Tranception is likely due to ProGen2 having almost 10 times as many parameters as Tranception, which may be compensating for its more limited training dataset. When controlling for model size, Tranception consistently outperforms ProGen2 (Fig. 2D-E, S5A,E).

In contrast to its strong performance on DMS datasets, SaProt performs substantially worse on real-world viral forecasting compared to other PLMs (Fig. 2F, 4F). Tranception was the top-performing model, both in identifying the most frequently occurring mutations (top panels, Fig. 2F, 4F, S6A) and in predicting emerging mutations (bottom panels). PLMs generally outperform DMS experiments for forecasting, with the exception of SARS-CoV-2 for which the available DMS assays measured only mutations that were already highly prevalent during the pandemic.

PLM performance on real-world evolution forecasting is complicated by data leakage (when test data is included in training). Because these models are trained on large datasets, many of the sequences used in our forecasting tests are likely present in their training data (Table S3). Consistent with this, Tranception’s accuracy declines for more recent clades, which are less likely to contain mutations seen during training (Fig. S6B). Nevertheless, in the most recent influenza clades (representing current H1N1 and H3N2 vaccine strains) where leakage is minimal, Tranception and ProGen2-XL outperformed the DMS experiment at forecasting (Fig. 2F, bottom).

#### Future directions for PLM improvements for viruses

Overall, our comprehensive comparison of modeling approach uncovers the drivers of successful viral modeling, and suggests that the low performance of existing PLMs on viral proteins is primarily driven by the poor representation of viral sequences in training data. While architectural improvements and increasing model size may help, these gains are ultimately constrained by the lack of viral training data. Increasing the breadth and diversity of viral sequences—particularly from underrepresented lineages—will be essential to build models that perform robustly across the viral protein universe. Models may benefit from training not only on unclustered datasets but also focusing on viral-specific databases including GISAID, Pathoplexus ^85,86^ and even the metagenome-derived VIRE ^87^, as well as relevant subsets of Uniref or larger databases like BFD, MGnify, and Logan ^88–90^. Although if the goal is not single-virus specific, care must be taken during sampling to avoid over-training on any one highly sequenced virus.

Incorporating structural information into PLMs may help mitigate the challenges posed by the sparse and highly divergent nature of viral sequence data. Structural features can evolve up to ten times slower than amino acid sequences ^91^ and some proteins, including viral proteins, retain structural similarity despite sharing less than 10% sequence identity ^92^ (Fig. S9A), enabling more reliable detection of distant homologs and improved protein function annotation ^93–95^. The benefits of structure-aware training is evident by SaProt’s higher performance on almost half of the EVEREST DMS benchmarks (including some fitness DMSs for relatively well represented viruses in the PDB) compared to sequence-only models (Fig. 2C,E, S5A). The inclusion of even a small number of viral structures found in the PDB yielded consistent improvement boost over SaProt-AF2 (which trained only on the AlphaFold2 database with no viral structures; Fig. S5C,D).

We find some evidence that structural training data may have a greater impact on performance than sequence data alone. For instance, SaProt outperforms ESM-1v despite being trained on orders of magnitude fewer amino acid data points (Fig. 2B, S9C). Second, the availability of additional training points yielded a larger improvement in performance for SaProt compared to Tranception (Fig. S9D). Additionally, SaProt’s performance is better predicted by the combined number of sequence and structure training points than by either alone (Fig. S9B). The growing availability of viral-specific structural databases ^94,96–98^, holds promise for improving viral modeling as structural information may provide local structural context or the ability to link relevant sequences with remote structural homology.

### Alignment-based Models have high and consistent performance on Viruses

#### Alignment relevance improves performance

Rather than learn from across the protein universe (and therefore primarily non-viral sequences), alignment-based models are trained on a set of homologous sequences and can therefore learn relevant evolutionary constraints for specific protein (Fig. 3A) ^2,3,18–20,30–36^. By focusing on homologous sequences, alignment-based models avoid pitfalls of PLM training datasets and perform equally well on viruses as other taxa (Fig. 3B), with even the simplest alignment-based model (site-independent PSSM) outperforming most PLMs for viruses (Fig. 2A).

**Figure 3:**
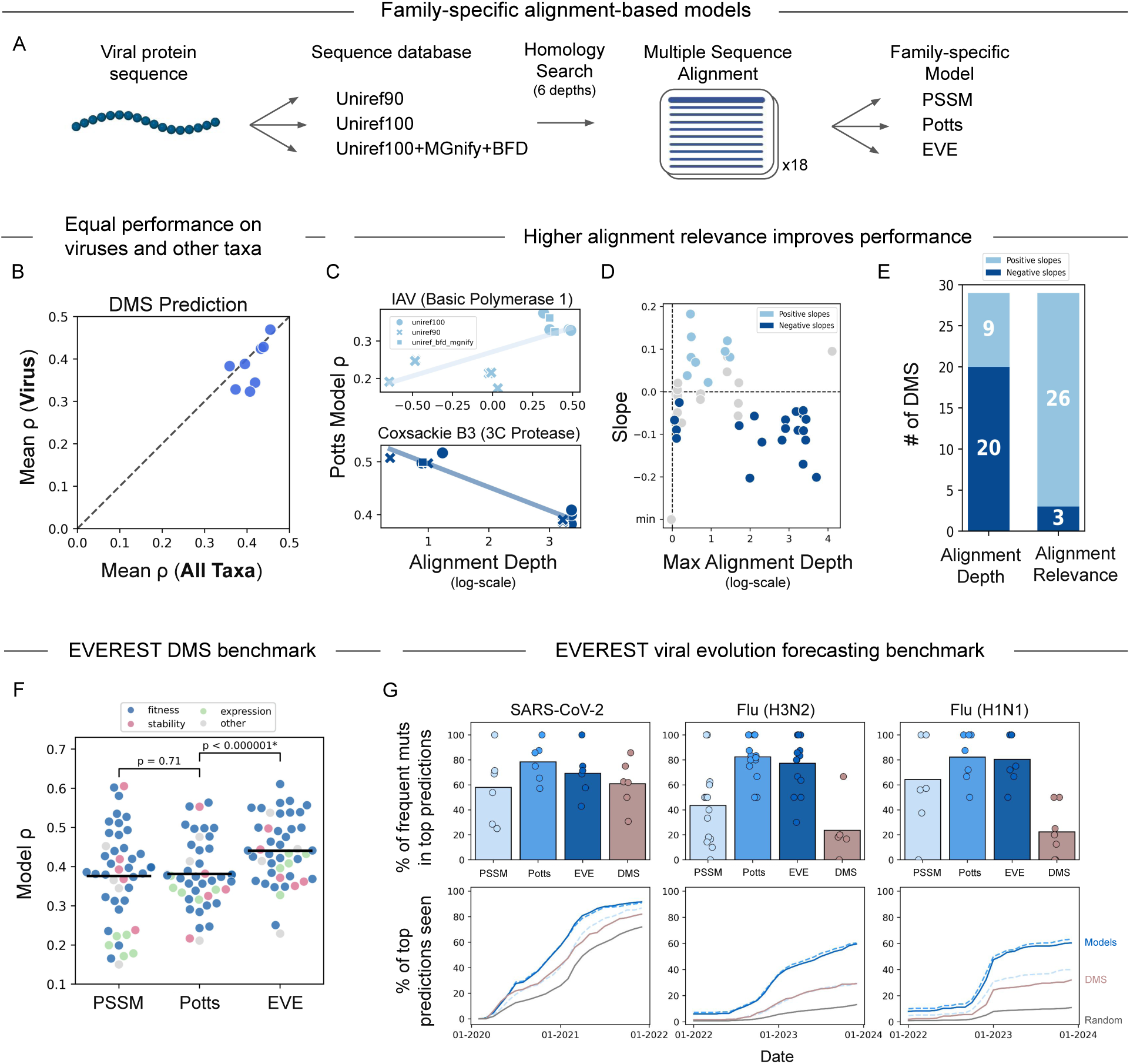
Evaluation of alignment-based models for viral evolution and mutation-effect prediction. **A.** Overview of MSA generation for alignment-based models. **B.** Unlike protein language models, alignment-based models have similar performance for viral proteins compared to proteins from other taxa. Performance is measured as Spearman correlation with experimental deep mutational scanning (DMS) assays from prior benchmark ^17^. **C.** Increasing alignment depth (Neff/L) improves Potts model ^18^ performance for influenza polymerase, but decreases performance for coxsackievirus protease. Performance measured as Spearman correlation between model prediction and DMS. **D.** Alignment depth is a poor predictor of Potts model performance. Colors indicate whether performance increased (light blue), decreased (dark blue), or was unchanged (gray; Spearman range *<* 0.1) with increasing alignment depth. **E.** Alignment relevance, measured as the fraction of sequences within 90% identity to the query sequence, is a better predictor of model performance than alignment depth. Reported is the number of viral proteins with positive slopes between Potts model performance and alignment-selection metric for DMSs where Spearman range is *>*0.1. **F.** Alignment-based VEP performance on EVEREST DMS benchmark. EVE ^19^ is a significant improvement over Potts and PSSM models (*P <* 0.001, paired t-test). Alignments selected using alignment relevance criteria. **G.** Alignment-based VEP performance on EVEREST viral forecasting benchmark. Top: percentage of highly frequent mutations captured by model predictions across 6 SARS-CoV-2 clades, 9 H1N1 clades and 15 H3N2 clades. Bottom: percentage of model-predicted mutations subsequently observed during evolution. Time-course analyses shown for representative clades—SARS-CoV-2 Wuhan (wild-type vaccine strain) and current flu vaccine strains for H1N1 (6B.1A.5a.2a.1), and H3N2 (3C.2a1b.2a2a.3a.1). Model predictions correspond to top 250 ranked mutations (see Fig. S7,S8 for alternate thresholds). Frequent mutations seen >1k times for flu and >10k times for SARS2.

A common challenge in using alignment-based methods is identifying which set of sequences will yield optimal model results in the absence of any evaluation data. Prior strategies for alignment selection often include selecting MSAs that recover structural contacts ^18^. However, most viral MSAs—except for a few notable cases such as HIV—fail to recapitulate structural contacts from couplings analysis (Fig. S10). In the absence of structural ground truth, a common heuristic is to instead maximize “alignment depth” (measured as the ratio of the effective sequence number to protein length, Neff/L), while ensuring high positional coverage ^18^. Neff represents the effective number of non-redundant sequences computed by down-weighting highly similar sequences. Alignment depth can be categorized as low (Neff/L*<*1), medium (1*<*Neff/L*<*100), or high (Neff/L*>*100) ^17^. To investigate the impact of alignment selection on model performance, for each protein in our EVEREST DMS benchmark we generated 18 MSAs by querying three sequence databases (UniRef90, UniRef100, and UniRef100+MGnify+BFD) and six depths for homology search (Fig. 3A).

Contrary to common practice, we find that deeper alignments do not necessarily improve performance (Fig. 3C-D). For example, while model performance for influenza A virus (IAV) polymerase improved with alignment depth (Fig. 3C, Top), deeper alignments yielded worse performance for Coxsackie B3 protease (Fig. 3C, Bottom). In fact, for most proteins in our EVEREST DMS benchmark, maximizing alignment depth reduced performance, particularly for proteins with the most sequences retrieved (Fig. 3D). We hypothesize that beyond a minimum alignment depth, incorporating additional sequences—especially from distantly related organisms—may introduce noise or conflicting constraints, as these sequences are subject to distinct selective pressures.

To test this, we examined whether performance correlated with “alignment relevance”, quantified as the proportion of MSA sequences sharing *>*90% identity with the target viral protein, given Neff/L *>*1. Alignment relevance is a stronger predictor of model performance than either alignment depth or the fraction of viral sequences in the alignment (Fig. 3E; S11A-B), though maximizing relevance also increases the fraction of viral sequences in the alignment (Fig. S11F). For both the coxsackievirus protease and IAV polymerase, selecting the alignment that maximized relevance improved Potts model performance (Fig. S11C,E).

Selecting alignments based on the relevance criteria significantly improved Potts model performance across the EVEREST DMS benchmark (from an average Spearman of 0.32 to 0.39; *P <* 0.001, paired t-test). Performance improvements were more pronounced for predicting fitness assays, where functional differences in divergent sequences seen in deeper alignments may be misleading, as opposed to stability or expression prediction where the learned phenotype may be more consistent across proteins from diverse species (Fig. S11G). Given these results, we propose a practical heuristic for alignment selection in viral proteins: (1) select alignments with at least medium depth (Neff/L *>*1), and (2) among these, choose the alignment with the highest fraction of sequences sharing *>*90% identity to the target. For proteins with no alignment reaching Neff/L *>*1 (only one protein in our dataset), we instead select the alignment that maximizes alignment depth.

Few of the alignments selected using relevance criteria required metagenomic sequences found in BFD or MGnify (Fig. S11D). This is likely because alignment relevance selects for closely related sequences while disfavoring the inclusion of highly divergent proteins found in BFD+MGnify ^99^. In some cases, these larger databases also caused memory issues during model training (Table S4), as BFD+MGnify is almost 9 times the size of Uniref100^89,100,101^. These results suggest that, for eukaryote-infecting viruses, expanding alignments to include metagenomic sequencing may rarely provide additional benefit, though this may change for more viral-focused datasets, or when focusing on non-fitness phenotypes.

#### EVE outperforms other alignment-based models on EVEREST

Using the alignment relevance selection strategy, we train three alignment-based VEPs with increasing epistatic complexity: a position-specific scoring matrix (PSSM; site-independent) ^18^, Potts model (an epistatic pairwise maximum entropy model) ^18^, and EVE (a fully connected variational autoencoder) ^19^.

We found that EVE consistently outperformed both PSSM and Potts models across nearly all EVEREST DMS datasets, and had a higher average Spearman correlation (*ρ* = 0.44 vs. 0.38 for both PSSM and Potts model; *P <* 0.0001, paired t-test; Fig. 3F, S13, Table S5). This is consistent with prior reports that capturing higher-order epistasis, using a variational autoencoder, improves prediction accuracy ^2,19,102^. Surprisingly given the frequent use of pairwise epistatic alignment-based models for viral fitness prediction ^20,30–36^, the Potts model and site-specific PSSM had comparable overall DMS performance for viruses, in contrast to other taxa (Table S2). This may be because pair-wise terms (inferred by the Potts model) are incorrectly learned in the limited diversity regime (consistent with poor structure reconstruction from coupling analyses for viral proteins; Fig. 2B, S10) and using alignment relevance advantages site-independent terms (by learning site-specific frequencies from a set of sequences with high similarity to the query). Supporting this, we find that PSSMs are more likely to outperform Potts models when the alignment is composed of more relevant sequences to the query (Fig. S12C).

Despite the comparable performance of PSSM and Potts models on the DMS benchmark (*P >* 0.5; paired t-test), we find that Potts models significantly outperform PSSM on forecasting viral evolution, with comparable accuracy to EVE models (Fig. 3G, 4F, S6A; *P <* 0.0001; paired t-tests). Differences in performance on the EVEREST DMS dataset and the viral forecasting dataset highlight that models that are better able to recapitulate specific phenotypes measured in laboratory setting are not necessarily better at capturing the full set of fitness constraints acting on evolving viruses, for which sequences from prior evolution may be particularly informative.

**Figure 4:**
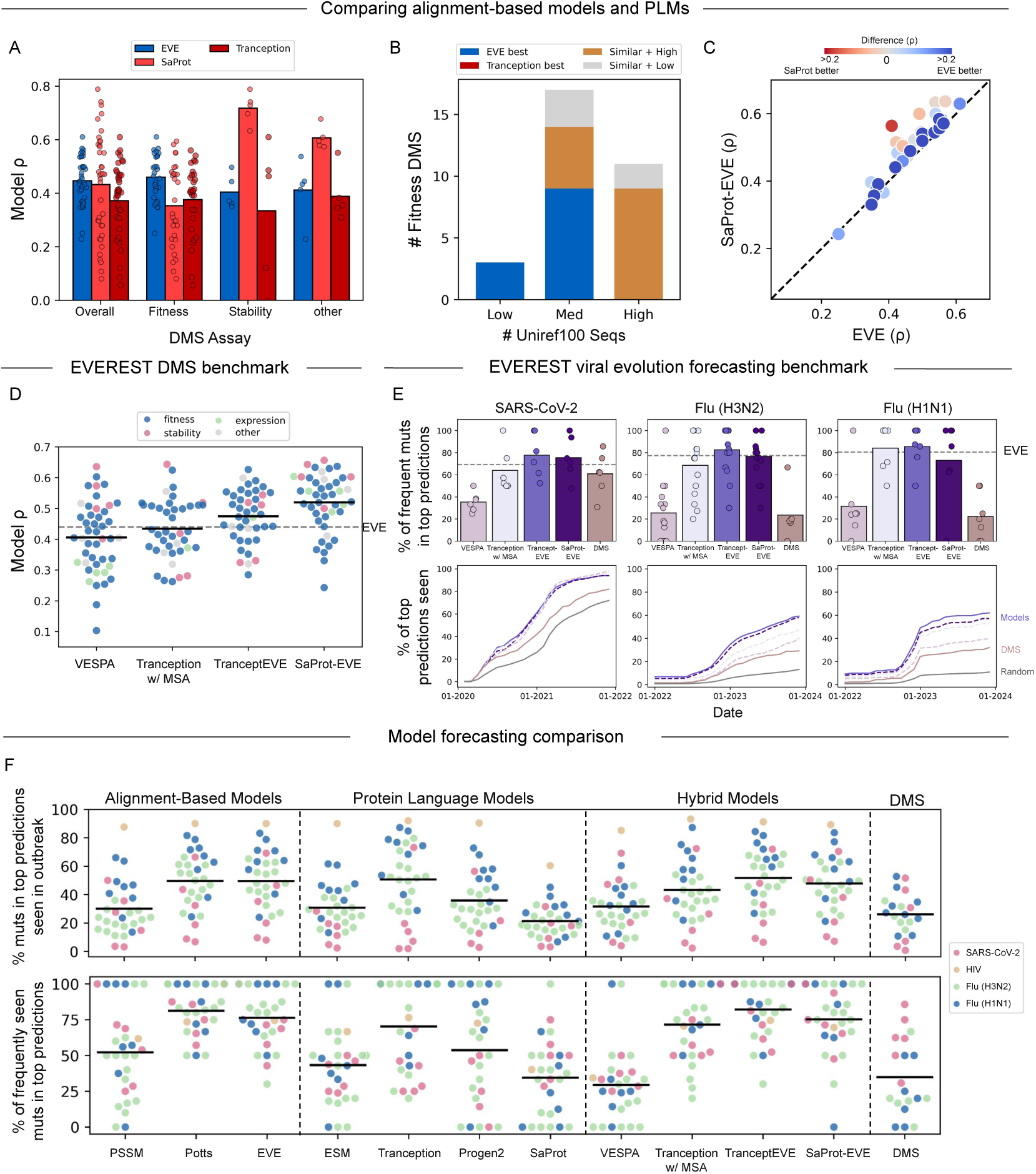
Combining PLM and alignment-based models improves performance. **A.** Alignment-based models and PLMs perform best on different DMS phenotypes. **B.** Relative performance of EVE and Tranception at different sequence-depth-regimes: Low (*<*3000), Medium (3000-30,000), High (*>*30,000). Colors indicate the best performing model (*ρ* difference *>* 0.1 between EVE and Tranception), or if the models had similar and high performance (*ρ* difference *<* 0.1, and *ρ >* 0.4) or similar and low performance (*ρ* difference *<* 0.1, and *ρ <* 0.4). This analysis considered fitness DMSs. Number of sequences determined by MMseqs search on UniRef100^104^ **C.** SaProt-EVE performs as well as EVE and exceeds EVE when SaProt is a better model. **D.** Hybrid models performance on EVEREST DMS benchmark. SaProt-EVE and TranceptEVE have improved performance relative to either model alone (EVE, dotted line), as well as other hybrid models (*P <* 0.005, paired t-test). **E.** Hybrid model performance on EVEREST viral forecasting benchmark. Top: percentage of highly frequent mutations captured by model predictions across 6 SARS-CoV-2 clades, 9 H1N1 clades and 15 H3N2 clades. Bottom: percentage of model-predicted mutations subsequently observed during evolution. Time-course analyses shown for representative clades—SARS-CoV-2 Wuhan (wild-type vaccine strain) and current flu vaccine strains for H1N1 (6B.1A.5a.2a.1), and H3N2 (3C.2a1b.2a2a.3a.1). Model predictions correspond to top 250 ranked mutations (see Fig. S7,S8) for alternative thresholds). Frequent mutations seen *>*1k times for flu and *>*10k times for SARS-CoV-2. **F.** Percentage of model-predicted mutations subsequently observed during evolution (top) and percentage of highly frequent mutations captured by model predictions (bottom) across 6 SARS-CoV-2 clades, 9 H1N1 clades, 15 H3N2 clades and 1 HIV clade.

### Comparing and combining alignment-based models and PLMs

#### Alignment-based model EVE outperforms PLMs at predicting viral mutation effects

We next compare the top-performing alignment-based variant effect predictor (EVE) and PLMs (Tranception and SaProt). Across the EVEREST DMS benchmark, EVE significantly outperformed Tranception (*P <* 0.01, paired t-test) and showed comparable performance to SaProt (*P >* 0.05, paired t-test; Fig. 4A). SaProt’s advantage was primarily driven by superior performance on stability assays, where it outperformed both EVE and Tranception (*P <* 0.01 and *P <* 0.05, respectively, paired t-tests). In contrast, for DMSs assessing general fitness, which better reflect selective pressures in naturally evolving strains, EVE outperformed both PLM models (*P <* 0.001, paired t-test; Fig. 4A, Table S5).

In contrast to the view of PLMs being better suited in low-sequence regimes ^44^, we find that EVE’s relative advantage over PLMs was particularly pronounced for proteins with few homologous sequences, as is likely in a new viral outbreak scenario (Fig. 4B, S14E). EVE’s success over PLMs must come not because the alignment has access to more data–as this comparison is with the Uniref100-trained Tranception model–but because of explicit access to the few related sequences that may become diluted by unrelated sequence families in a PLM. While both Tranception and SaProt showed an improvement in performance with the number of relevant training points (Fig. S9C-D), EVE’s performance was consistent across different number of related sequences (Fig. S9D, S12A, S14E)—for example, while EVE had high performance for well-sequenced viruses, like HIV and Flu antigens (*ρ >* 0.4), they were far from the best performing proteins, which are primarily non-antigens (Fig. S12A, S14C-D).

The contrasting observations that PLMs benefit from training on larger datasets with more remote homologs while alignment-based models perform better with more focused alignments, likely stems from the fact that alignment-based models (besides redundancy-weighting) treat all sequences identically regardless of sequence and functional similarity to the target. In contrast, PLMs, by learning across the protein universe, must by necessity learn to reason about sequences with varying degrees of relevance.

The superior performance of EVE on fitness DMS assays would suggest that it is better suited for predicting real-world viral evolution. While Tranception and EVE have similar performance in forecasting ( *P >* 0.5; paired t-tests; Fig. 4F), this is likely due to data leakage where sequences in the test set are seen during Tranception training. Consistent with this, the apparent advantage of Tranception over EVE is most pronounced for clades that emerged earlier (and were included in training), but diminishes for the most recently circulating clades, which were not part of the training data (Fig. S6B). Moreover, while SaProt excels at predicting stability, only a subset of stable proteins satisfy other necessary fitness constraints, such as receptor binding, conformational changes, or adaptive selection pressures captured by evolutionary sequences. This difference is evident when comparing the distribution of top scores for SaProt and EVE—many mutations receive equivalently high SaProt scores, while only a small fraction achieve the highest EVE scores (Fig. S18). High-frequency mutations that emerge during evolution tend to require both high EVE and high SaProt scores, but EVE’s more selective scores makes it a better discriminator of evolutionary trajectories.

#### Hybrid models improve performance over alignment-based or PLMs individually

Several strategies have been proposed to combine alignment-based models with PLMs. One approach is to fine-tune PLMs using sets of unlabeled homologous sequences (“evotuning”) ^103^. Another is to construct hybrid models that combine mutation scores between both approaches. We examined multiple hybrid strategies, including Tranception with site-independent MSA retrieval ^22^, TranceptEVE (Tranception-EVE hybrid) ^38^, a newly constructed SaProt-EVE hybrid, and VESPA ^79^, which integrates PLM embeddings with conservation and BLOSUM substitution scores. Ideally, a hybrid model should perform at least as well as its best component and, where possible, surpass it through synergistic integration.

While evotuning did improve PLM performance (SaProt *ρ* = 0.28 vs. best evotuning *ρ* = 0.34 for Lassa DMS; Methods), it did not reach the performance of even a site-independent alignment-based model (*ρ* = 0.51), indicating that simple fine-tuning is not always sufficient to surpass alignment-based approaches. These observations are preliminary, and further evaluation at scale is warranted.

Alternatively, we find that hybridizing alignment-based models with PLMs (e.g., SaProt-EVE and TranceptEVE) yields superior performance relative to either modeling approach alone (Fig. 4C-D, S14A,S15B-C). In fact, SaProt-EVE and TranceptEVE significantly outperformed all other models—including alignment-based models, PLMs, and alternative hybrids—on the EVEREST DMS benchmark (*P <* 0.005, paired t-test; Fig. 4D, S15D). However, the benefits of hybrid approaches for viral forecasting were more modest (Fig. 4E-F), even while plagued with the same data leakage concerns as PLMs which should inflate performance. Overall, they represent a robust default strategy when experimental validation is unavailable, multiple phenotypes are of interest, or a single, reliable model is preferred over multi-model evaluation.

### Calibrating Reliability Estimates

We have thus far identified conditions under which different modeling approaches perform best. What remains unclear is how to know *a priori*, in the absence of experimental data, how reliable model predictions are for a specific viral protein. Reliability estimation has been useful in other biological domains, most notably for AlphaFold2^105^, but such estimates have not yet accompanied variant effect predictions. To address this, we calibrate reliability metrics on our EVEREST DMS benchmark tailored to each modeling approach.

For alignment-based VEPs, we find that the number of effective sequences within 90% identity (Neff @ 90% ID) is the best predictor of model reliability (Fig. 5A, S12A). For most PLMs, we find that pseudo-perplexity (PPPL), a measure of model uncertainty given a specific input ^25,106^, correlates with model performance across proteins (Fig. 5B,S16A). We scaled PPPL to a value between 0 and 1, where higher values reflect greater confidence. This metric was a strong predictor of SaProt performance (*ρ* = 0.75), outperforming the number of homologous training sequences and structures (*ρ* = 0.68; Fig. S9). For some models other metrics were more predictive of performance—for example, Tranception performance was better explained by number of relevant training data points (Fig 9D).

**Figure 5:**
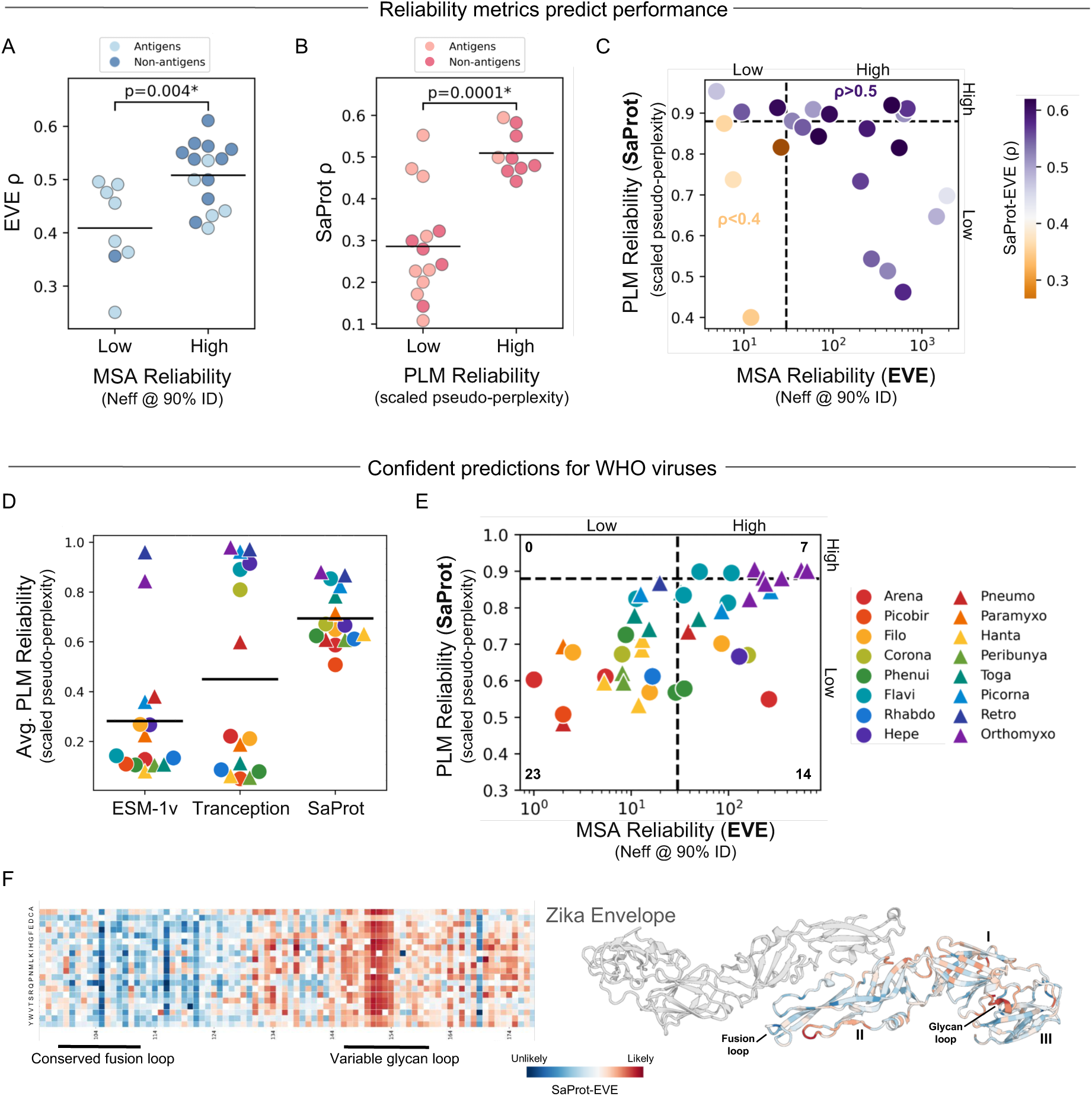
Quantifying model reliability across WHO priority viruses. **A.** EVE models with high MSA reliability scores have significantly better performance than those with low reliability. MSA reliability is measured as the number of effective sequences (Neff) with greater than 90% identity to the query (Neff @ 90% ID). **B.** SaProt models with high PLM reliability scores have significantly better performance than those with low reliability. PLM reliability is measured as a scaled pseudo-perplexity. **C.** Hybrid model SaProt-EVE has correlation below 0.4 only where both reliability metrics for EVE and SaProt are low and correlation greater than 0.5 when both reliability metrics are high. **D.** Inferring PLM reliability for 44 viral antigens from 40 WHO-prioritized viruses. Structure-aware PLM, SaProt, has on average higher reliability than sequence-only PLMs, ESM-1v and Tranception. Plotted are the average reliability score for a given viral family. **E.** EVE and SaProt reliability scores for all WHO viral antigens. More than half of WHO antigens have both low MSA and PLM confidence. Based on our viral benchmarks, antigens with high model confidence are expected to yield Spearman correlations greater than 0.4 with DMS experiments. **F.** SaProt-EVE predictions for priority virus Zika envelope, showing *in silico* mutational scan and site-average fitness scores mapped on the structure (AlphaFold3^108^ structure and PDB 5ire). Fusion loop (98-107) and glycan 150 loop (147-161).

For each of the MSA and PLM reliability metrics, we identify stringent thresholds for high versus low model reliability based on maximizing sensitivity and specificity (Youden’s J) for predicting high model performance (defined as Spearman *ρ >* 0.4). We find that for EVE, a MSA reliability metric of 30 effective sequences within 90% identity effectively separates high versus low performance (*P <* 0.01; t-test). For SaProt, a scaled PPPL of 0.88 (interpretable as an average of ≈3 amino acid choices per residue) similarly separates high performance (*P <* 0.0001; t-test). Importantly, proteins exceeding the high confidence threshold consistently have *ρ >* 0.4, indicating that these cutoffs are conservative estimates.

The MSA and PLM reliability scores also predicted which of the two modeling approaches would perform best for a given viral protein (Fig. S14B). Notably, across fitness assays, non-antigens have consistently higher MSA reliability scores (likely as non-antigens are under less antigenic pressure and therefore have more homologs within 90% identity; Fig. 5A, S14C-D). However, many of the highest PLM reliability scores are for common antigens in the PDB, despite potential for the greater mutability of antigens being interpreted as greater model uncertainty (Fig. 5B, S14C-D).

When applied to the EVEREST DMS dataset, hybrid models with multiple reliability metrics can work in tandem–SaProt-EVE only underperforms (Spearman *ρ <* 0.4) when both reliability scores fall below their respective thresholds. Conversely, all viral DMS experiments with both reliability scores above threshold exhibit correlations exceeding 0.5 (Fig. 5C), demonstrating that high-confidence predictions across both modeling approaches provide synergistic estimates of variant effects representative of both the model’s internal confidence in its predictions and the richness of evolutionary information supporting those predictions.

#### Future directions

While EVE and SaProt reliability thresholding significantly separate DMSs by performance, the MSA reliability metric is less accurate than the PLM metric. PPPL is an intrinsic measure of a model’s predictive uncertainty, and is a property of the PLM predictions itself rather than the input data (which is the case for EVE). We find that an equivalent “pseudo-perplexity” measure for EVE does not distinguish high versus low performance, this is likely due to EVE’s consistent performance in the EVEREST DMS benchmark so there is less signal to separate high versus low performance (Fig. S16C). Future work evaluating this and other intrinsic alignment-based model reliability metrics is warranted.

### Evaluating 40 WHO Priority Viral Pathogens

The WHO recently prioritized 40 RNA viruses (with 44 antigens as some viruses have multiple antigenic proteins) for research and countermeasure development, based on their potential for transmission, virulence, and the current lack of effective medical interventions ^1^. This list includes both extensively studied pathogens—such as influenza, HIV, and SARS-CoV-2—and less characterized viruses like Lassa, Nipah, and Lujo viruses ^107^. Leveraging our calibrated reliability metrics, we aimed to identify the most effective modeling approaches for antigenic proteins from each WHO-prioritized virus, particularly those where scarce experimental data limits direct model evaluation. We assessed whether current models provide sufficient accuracy or whether improved models and additional data are needed to enhance predictions.

Using our calibrated metrics, we find that existing state-of-the-art models will likely fail to reliably predict the functional impacts of mutations in a large fraction of these antigens (Fig. 5E; Table S6). Of the 44 WHO viral antigens, only 21 have sufficient sequence diversity to be accurately modeled using EVE. Only 7 of the 44 antigens—mostly influenza hemagglutinins—demonstrated high PLM confidence for SaProt, the PLM with greatest average reliability (Fig. 5D). EVE’s low reliability may seem surprising given its strong performance on DMS benchmarks, but these benchmarks likely overestimate model generalizability due to their focus on well-characterized viruses (Fig. 17C). Applying reliability estimates to WHO-prioritized viruses therefore provides a more balanced assessment of current model limitations.

These findings highlight that additional sequences and structures are required for more accurate modeling of pandemic risk pathogens. Many under-sequenced viral families and structural classes cluster together with low reliability scores (Fig. 5E, S17A-B). For instance, most Class II fusion accompanying proteins show low reliability and have some of the lowest numbers of related sequences in UniRef (both pre- and post-clustering), reflecting biases in available training data and structural coverage. Given these limitations, aggregating scores from the best-performing models is expected to mitigate some of these issues. Moreover, even “unreliable” predictions from understudied viruses may still prove useful for prioritizing mutations or identifying conserved regions in the absence of experimental data. We provide model predictions for all 44 antigens across the 40 prioritized viruses, alongside estimated model reliability scores, as a community resource.

We anticipate these predictions can be used in multiple ways, given that most of these viruses lack extensive experimental characterization. For therapeutic design, it is essential to identify evolutionarily conserved regions less likely to mutate in the future. For example, for the Zika envelope protein dimer, a class II fusion protein, model predictions reflect the known evolutionary constrained fusion loop and variable glycan loop ^109^ (Fig. 5F). For newly emerging viruses, top mutation predictions could also be incorporated into therapeutic designs. As an example of the predictive power of these models, we highlight the H1N1 “swine flu” pandemic—a novel spillover from pigs distinct from prior flu pandemics—and the predictability of the S179N mutation that necessitated the first vaccine strain update ^110^. EVE, without any training leakage, predicted this mutation in the top scores of all three clade models prior to the update. Evolutionary sequence models therefore identified this major immunogenic change at the very start of the H1N1 pandemic, six years before its eventual dominance.

### Biosecurity Assessment

Biosecurity requires a careful balancing of risks and benefits, which are inherently coupled. Improvements in predictive modeling can substantially benefit public health, while simultaneously increasing the potential for misuse. The same capabilities that enable rational vaccine and therapeutic design could, in principle, be repurposed to engineer more virulent or immune-evasive pathogens.

To limit the potential for misuse of the EVEREST dataset, we deliberately assess only a narrow and constrained component of the broader viral design problem relevant to biosecurity. Our analyses are limited to models that predict the effects of single amino-acid substitutions in individual viral proteins. We exclude nucleotide-level effects, combinations of mutations, insertions and deletions, and full viral genome contexts. We further restrict our benchmarks to functions relevant to viral surveillance and therapeutic design, rather than explicitly harm-enhancing phenotypes such as increased pathogenicity or virulence, and we exclude viral antibody-escape deep mutational scanning datasets. In addition, we focus on functions learnable from naturally occurring evolutionary sequences and structures, rather than those relevant under misuse-oriented conditions (e.g., non-physiological temperature or pH ranges).

To enable pandemic preparedness, we show for the first time that current mutation effect predictors are unreliable for more than half of pandemic-threat viruses. While this hinders the immediate biosecurity risk of these models generalizing to a wide range of viruses with potential for harm, it also highlights that limits of current models. Nevertheless, viral performance has improved over time (Fig. S16B), and is likely continue to improve with better models, more viral sequences and newly available structure data ^94,96–98^, and finetuning on specific viruses ^46,111–113^. In this context, EVEREST provides not only a framework for evaluating models relevant to pandemic preparedness, but also a mechanism for monitoring the emergence of dual-use biosecurity risks as predictive capabilities advance.

## Conclusion

In this study, we present the first virus-specific, large-scale comparison of variant effect predictors, evaluating alignment-based models, protein language models (PLMs), and hybrid approaches across 45 deep mutational scanning experiments and 31 real-world forecasting datasets using our EVEREST benchmark. We also introduce calibrated reliability metrics that quantify model confidence in the absence of validation data—a scenario typical of emerging viral outbreaks. Applying these metrics reveals that only approximately half of the 40 WHO priority RNA viruses can currently be predicted with high reliability, underscoring fundamental limitations in existing methods and data for many under-studied pandemic threats.

Alignment-based models trained on family-specific viral sequences frequently outperform PLMs trained on large, general protein corpora. Notably, this advantage persists even in low-sequence-depth regimes characteristic of novel outbreaks, where “single-sequence” PLMs were previously assumed to yield superior performance. These results suggest that widely used PLMs such as ESM-2 are often ill-suited for viral mutation effect prediction, despite their growing use in this domain ^44–46^.

The relative underperformance of PLMs on viruses likely reflects their limited representation in training data, which is disproportionately reduced in clustered sequence datasets (e.g., UniRef90 and UniRef50). Whereas PLMs benefit from scale and diversity across distant protein families in training, alignment-based approaches benefit from training on more evolutionarily relevant sequences with higher identity to the query. This contrast highlights a fundamental modeling difference: alignment-based methods effectively treat all homologous sequences as equally informative (after redundancy weighting), while PLMs implicitly attempt to reason over relevance—a task made difficult when viral sequences are sparse or underrepresented.

These findings suggest several priorities for future research. First, PLM performance on viral proteins may be improved through training on larger, unclustered datasets, viral-specific pretraining or fine-tuning, domain adaptation, or the integration of additional data modalities ^46,111–113^. Structure-aware modeling, in particular, may help compensate for sparse viral sequence data by leveraging more conserved structural constraints. Similar strategies may be necessary for RNA language models applied to viruses, which likely suffer from analogous dataset limitations ^114^. Second, alignment-based models could be strengthened by weighting sequences according to evolutionary proximity to the query, rather than relying solely on clustering heuristics, and by developing intrinsic, model-based reliability estimates. Third, alternative approaches also offer promising ways of improving viral mutation effect predictions. For instance, GEMME—a phylogeny-based model incorporating mutation frequency, evolutionary accessibility, and sub-tree conservation—achieves performance comparable to the best models in our benchmarks (fitness DMS prediction, *ρ* = 0.5; Table S5). Fourth, evaluation strategies should extend beyond single-mutation concordance, incorporating multi-mutation genotypes and strain-level effects that better reflect real viral evolution.

Many of the guidelines for future model improvements outlined here for viruses are likely to generalize to other proteins with limited sequence diversity. Yet, for many pandemic-risk viruses, with limited existing data, methodological advances alone are unlikely to suffice. Increased sequencing efforts, not only on the WHO priority viral species themselves, but on related viruses in their natural reservoirs, will be essential for improving modeling and enabling pandemic readiness. Notably, SARS-CoV-2 modeling success stems not from the millions of highly similar COVID-19 pandemic sequences, but from approximately 3,000 diverse pre-pandemic coronavirus sequences sampled primarily from bats and pigs, which provided the essential evolutionary and functional template for rapid pandemic response ^2^.

## Data and Code Availability

Data, results, and code are available at https://github.com/debbiemarkslab/priority-viruses for 45 viral DMSs, 44 WHO-priority viral antigens, and viral forecasting of 31 clades. Code for model training and for mutation effect scoring is available through ProteinGym ^17^ at https://github.com/OATML-Markslab/ProteinGym. DMS data used for model evaluation collected and standardized from papers listed in Table S1^4,50,54–78^. Sequence data analyzed for forecasting from GISAID and LANL ^85,115^.

## Acknowledgments

The authors thank Aaron Kollasch and other members of the Marks lab for valuable discussions. Special thanks to the teams of experimentalists who developed and performed the viral DMS assays this work is built on. If you are using these assays in your work, please cite the corresponding papers in Table S1. This work was supported by the Coalition for Epidemic Preparedness Innovations (CEPI), including Danny Scarponi, and biosecurity analysis completed alongside experts from CEPI, particularly Andrew Hebbeler (Biosecurity Director). S.G. was supported by funding from the FutureHouse AI-for-Science Postdoctoral Fellowship; work at FutureHouse is supported by the generosity of Eric and Wendy Schmidt. We gratefully acknowledge all data contributors, i.e., the authors and their originating laboratories responsible for obtaining the specimens and their submitting laboratories for generating the genetic sequence and metadata and sharing via the GISAID Initiative, as well as LANL, on which some of this research is based.

## A Methods

### A.1 EVEREST Deep Mutation Scanning Benchmark

We curated and standardized a collection of 45 viral deep mutational scanning (DMS) datasets from 25 publications ^4,50,54–78^, more than doubling the number of viral datasets included in previous general-purpose benchmark (ProteinGym ^17^). The resulting EVEREST DMS benchmark spans viruses of major importance for vaccine development—including SARS-CoV-2 and other sarbecoviruses, seasonal and pandemic influenza viruses, and high-priority pathogens such as Lassa and Nipah—as well as viruses relevant to viral vector design (e.g., AAV).

We systematically searched for all available viral DMS experiments and included datasets measuring single–amino acid substitutions and phenotypes directly relevant to pandemic preparedness, such as protein expression, host receptor binding, viral replication, and infectivity. Datasets were excluded when the assayed phenotype was not directly comparable across viruses or was not aligned with the goals of this benchmark. Specifically, we excluded assays measuring drug inhibition or viral escape (from antibodies or polyclonal sera). While relevant for pandemic preparedness, viral escape datasets were excluded due to the limited number of antibodies or sera tested for most viruses, which can result in apparent false positives where a model correctly predicts escape but the DMS assay does not capture it.

There are some limitations in using DMSs as a benchmark. DMS phenotype single proteins under laboratory conditions that incompletely capture the full constraints a virus faces in the real world. DMSs also measure a narrow fitness landscape of largely single mutations. Yet, for comparing model performance broadly across diverse viral families, a new viral-specific DMS dataset remains our best available benchmark.

### A.2 EVEREST Viral Forecasting Benchmark

We created a new benchmark for evaluating the ability to predict viral evolution across multiple viruses and numerous phylogenetic clades: 9 influenza (H1N1), 15 influenza (H3N2), 1 HIV, and 6 SARS-CoV-2. Extensive surveillance sequencing allows us to test the ability of evolutionary sequence models to forecast frequent mutations that emerge in each clade in a retrospective study. We focus on single nucleotide mutations since multi-nucleotide mutations are very rare (i.e., *<* 4% of the test dataset for influenza, and *<* 1% for SARS-CoV-2). For alignment-based models, we perform strict time separation between training and testing datasets, while for PLMs we acknowledge and assess data leakage (test mutations included within PLM training data) that likely artificially inflate performance.

#### A.2.1 INFLUENZA H1N1 AND H3N2

##### Clade definitions

We perform phylogenetic analysis of H1N1 and H3N2 clades using the annual and interim reports sent from the Worldwide Influenza Centre at the Francis Crick Institute to the WHO to inform vaccine composition ^116^. Tracking the evolution of H1N1 and H3N2 from 2009 using these reports, we defined each emerging clade’s characteristic mutations, identified its most basal strain, and noted which years it was dominant. We used the basal strain as reference sequences for modeling each clade, reverting any unique adaptations (e.g., egg-passaged Q240R) to ensure that the reference only contained the clade’s characteristic mutations. We examined the following nested clade lineages for H1N1: pdm09, 6, 6B, 6B.1, 6B.1A, 6B.1A.5a, 6B.1A.5a.2a, and 6B.1A.5a.2a.1. We examine the following clade lineages for H3N2: 3C, 3C.2, 3C.2a, 3C.2a.1, 3C.2a2, 3C.2a.1b.1, 3C.2a.1b.2, 3C.2a.1b.2a.2, 3C.2a.1b.2a.2a.3, 3C.2a.1b.2a.2b, 3C.2a.1b.2a.2a.3a.1, 3C.3, 3C.3a, and 3C.3a.1.

##### Training Datasets

We assembled a training dataset of all full-length influenza A hemagglutinin protein sequences from the Global Initiative on Sharing All Influenza Data (GISAID) database ^85^, comprising 365,500 full-length sequences submitted before January 1, 2025. To generate an MSA per clade for alignment-based models, we filtered the full Influenza A dataset to include only sequences collected before that clade’s emergence, then deduplicated and aligned them to the identified clade reference.

##### Testing Datasets

Each model was evaluated on a corresponding test dataset containing all human sequences collected during the 2-4 year period for which that clade was dominant and annotated as belonging to that clade or its derivative subclades. For instance, sequences labeled 6B.1 or 6B.1A were included in the test set for the clade 6B model. We assigned clade labels based on our identified characteristic mutations, requiring only the new mutations specific to the child clade rather than cumulative acquisition of all ancestral mutations. To ensure this assumption would not mischaracterize strains, we verified the majority of strains assigned to a given clade contained at least 80% of the clade’s ancestral mutations. For forecasting analyses, we restricted to mutations observed at least 3 times, and classified mutations as “high-frequency” if they were observed more than 1,000 times.

#### A.2.2 HIV

##### Training Dataset

A set of HIV-1 subtype-stratified multiple sequence alignments were sourced from Lewitus et al. (2024) ^117^, in which HIV sequences from Los Alamos National Laboratory (LANL) were filtered and stratified to create complete and independent sequence sets that capture the diversity of each subtype phylogeny ^115^. All non-subtype C envelope (Env) protein alignments were concatenated and aligned to a reference sequence using MAFFT ^118^, creating an alignment of 3,452 effective sequences ^118^. This reference sequence was sourced from the LANL database as one of the earliest published subtype C Env sequences which was recorded in the 1980s ^119^.

##### Testing Dataset

All subtype C Env sequences available in the LANL HIV Sequence Database ^115^ were downloaded and aligned to the 1980 reference sequence ^119^ using MAFFT ^118^. Analysis was stratified to all sequences in the years 1986-1995 to create a time period more comparable to other virus in the EVEREST forecasting benchmark, with an end date chosen for when the observed fraction of mutations began to plateau, although this is still several times longer than the other test periods choices. This test split is phylogenetic rather than temporal (i.e., training data even for alignment based models includes sequences after this testing period). For forecasting analyses, we restricted to mutations observed at least 100 times, and classified mutations as “high-frequency” if they were observed more than 10,000 times (due to increased sequencing).

#### A.2.3 SARS-COV-2

##### Training Dataset

We selected five SARS-CoV-2 variants which were ancestors to the majority of circulating strains during the COVID-19 pandemic: Wuhan, BA.1, BA.4, BA.5, XBB, and JN.1. For each variant, we constructed a training alignment consisting of (i) all homologous Spike protein sequences available prior to 2020 (∼ 3,000 sequences), and (ii) all SARS-CoV-2 Spike sequences deposited in GISAID ^85^ prior to the emergence of that specific variant (Fig. S1D). Variant emergence dates were obtained from NextStrain ^120^.

##### Testing Dataset

For each parent variant, we used NextStrain ^120^ to identify all descendant lineages. We then subset GISAID to include only sequences sampled after the variant’s emergence and assigned to its descendant lineages. From these sequences, we extracted all single-nucleotide mutations observed within the two years following the variant’s first appearance. For forecasting analyses, we restricted to mutations observed at least 100 times, and classified mutations as “high-frequency” if they were observed more than 10,000 times (due to increased sequencing).

### A.3 Sequence and structure datasets

Both alignment-based and protein language models rely on sequence and structure databases either for retrieving homologous protein sequences or for PLM training.

UniRef100, is a database of a non-redundant protein sequence database containing all UniProt Knowledgebase records plus selected UniParc records ^101^. UniRef90 and UniRef50, are 90% and 50% identity-clustered versions of UniRef100. The Big Fantastic Database (BFD) integrates sequences from UniProt (Swiss-Prot and TrEMBL) ^101^, Metaclust ^88^, and the Soil and Marine Eukaryotic Reference Catalog ^100^. MGnify includes proteins from metagenomic assemblies ^89^.

AlphaFold databse (AF2DB) is comprised of 40 million AlphaFold predicted structures for UniRef50 sequences, which notably excludes all viral proteins. Open MetaGenomic (OMG) corpus (50% ID clustering) from combining IMG ^121^ and MGnify contains 200 million representative sequences from clusters with at least two members, a more 3-fold increase in sequence diversity compared to UniRef50^122^.

### A.4 Alignment-based models

#### A.4.1 Generation of multiple sequence alignments

All alignment-based models rely on a method for generating a multiple sequence alignment on which they are trained. Multiple sequence alignments of the corresponding protein family were obtained using the method outlined in Hopf et al. (2017) ^18^. This involved five search iterations of the profile HMM homology search tool JackHMMER ^123^ against the specified sequences database. We evaluated the impact of searching against three database with vastly different number of sequences: UniRef100, UniRef90, and UniRef100+BFD+MGnify.

We generated alignments with six different length-normalized bit scores: 0.5, 0.3, 0.1, 0.05, 0.03, and 0.01 bits per residue. A challenge during the process of fitting alignment-based models was the memory required to query large sequence databases. Some models encountered memory limitations that prevented training on the largest alignments (Table S4), but these would mostly be excluded by the choice of Neff cutoff. This highlights the computational burden of alignment-based approaches, particularly when using comprehensive databases like BFD and MGnify, with 2.1 billion and 2.4 billion sequences, respectively.

The alignments were post-processed to exclude positions with more than 50% gaps and to exclude sequence fragments that align to less than 50% of the length of the target sequence. We slightly relax the constraint on minimum column coverage for sequences in the training MSAs (50% instead of the 70% in Frazer et al. (2021) ^19^) as it led to superior fitness prediction performance in our hyperparameter tuning analyses for the different viruses modeled in our prior work ^2^. Similarly, following prior work, we also cluster sequences at 99% sequence identity (referred to as theta) instead of 80% sequence identity ^2,19,102^ due to limited viral sequence diversity and an expectation that viral sequence are more likely to contain constraint information within smaller differences than are proteins from higher-order organisms. For the best performing model EVE, where clustering is used strictly for sampling, there is no difference in performance however between clustering differences (Fig. S12B). See Reliability Estimation below for more details on sequence clustering and alignment selection.

#### A.4.2 PSSM

Position-specific scoring matrix (PSSM) models assume each position in the protein evolves independently and assigns a prediction score for each mutation dependent on its frequency in the alignment. We used the site-wise maximum entropy model as implemented in Hopf et al. (2017) ^18^.

#### A.4.3 Potts Model

To predict the effects of mutations that explicitly captures pairwise residue dependencies between positions, we used EVmutation as implemented in Hopf et al. (2017) ^18^. In a Potts model, protein families are represented as probability distributions governed by two constraint types: individual position preferences for specific amino acids/nucleotides and pairwise coupling constraints between positions. This approach assumes that complex correlations in sequence alignments arise from simpler underlying positional couplings, which traditional metrics like mutual information cannot properly disentangle due to transitive correlations. The resulting distribution represents the maximum entropy solution—the least-structured distribution that still matches the observed single-site and pairwise frequencies in the alignment.

#### A.4.4 EVE

To predict the effects of mutations capturing high-order dependencies between positions, we used EVE, a Bayesian Variational Autoencoder, as implemented in Frazer et al. (2021) ^19^. The architecture consists of a symmetric encoder and decoder architecture, each with 3 layers with 2,000-1,000-300 and 300-1,000-2,000 units respectively, as well as a 50-dimensional latent space. As generative models, VAEs can learn a complex distribution of the high-dimensional data on which they are trained, in our case, sequences from a specific protein family. We use single EVE models, rather than an ensemble of independent models as was reported in Thadani et al. (2023) ^2^. Note, that we use the negative of the evolutionary index reported by the model.

#### A.4.5 Imputing missing data

For accurate comparisons across the same sets of mutations in DMSs, we impute missing model values (for positions with insufficient alignment coverage, using a lenient coverage threshold) for alignment-based models with the mean value across the target protein. All mutation scores are calculated by PLMs.

### A.5 Protein language models

#### A.5.1 Tranception

Tranception ^22^ combines an autoregressive protein language model with inference-time retrieval from a MSA. We used Tranception Large (700M parameters) trained on UniRef100 using only the autoregressive inference without MSA retrieval as implemented in ProteinGym ^17^. Tranception averages the log ratios from both left-to-right and right-to-left scoring.

#### A.5.2 ESM-1V

ESM-1v ^21^ has a Transformer encoder architecture similar to BERT [Devlin et al., 2019] and was trained with a Masked-Language Modeling (MLM) objective on UniRef90. We use the implementation presented in ProteinGym ^17^ to handle sequences that are longer than the model context window (i.e., 1023 amino acids), and ensemble across 5 models.

#### A.5.3 PROGEN2

ProGen2^24^ is an autoregressive transformer with left-to-right causal masking. We primarily use ProGen2-XL trained (6.4B parameters) trained on both Uniref90 and BFD30.

#### A.5.4 SAPROT

SaProt ^41^ introduces a structure-aware vocabulary, into protein language modeling by training on Foldseek ^124^ 3Di tokens which represent the local geometric conformation information of each residue relative to its spatial neighbors. These 3Di tokens are combined with typical amino acid residue tokens as input to the SaProt model, which utilizes an ESM-2 Transformer architecture ^25^ but expands the embedding layer to encompasses 441 structurally-aware tokens instead of the original 20 amino acid residue tokens. We use (1) SaProt-650M-AF2, trained on approximately 40 million AF2 sequences/structures (from UniRef50) which notably explicitly excludes all viral proteins though implicitly included hundreds of thousands of prophages; (2)SaProt-650M-PDB, which continuously pre-trains the SaProt-650M-AF2 model on the PDB; and (3) SaProt-1.3B-AFDB-OMG-NCBI with 40 million AF2 structures, and 200 million OMG (50% identity filtering) and 150 million NCBI (70% identity filtering) sequences trained with multimodal input integration.

For structure inputs to Foldseek calculation, we fold monomeric forms of each DMS or WHO viral protein with AlphaFold3, where structures had not been folded previously in ProteinGym.

### A.6 Mutation effect scoring

Generative models learn from the distribution of protein sequences collected as a result of billions of evolutionary experiments to capture the biochemical and structural constraints governing functional proteins. These models are trained to learn the distribution of natural, functional sequences.

For a given protein *x* composed of residues (*x*_1_*, x*_2_*, …, x_L_*) the relative fitness of mutated protein compared to its wild-type can be calculated in the following ways depending on the modeling objective.

The fitness of a mutant sequence *x*^mutant^ is calculated as:

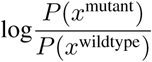

For ProGen2^22^, an autoregressive model, the likelihood of *x* factorizes via the chain rule and is calculated as:

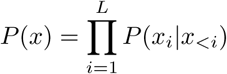

For Tranception, an autoregressive model with bidirectional scoring:

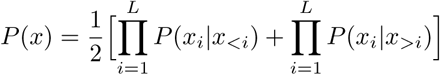

In the masked language model setting, for ESM-1v ^21^ and SaProt ^41^, we use the masked marginal scoring function instead:

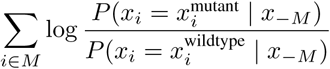

where *x*_−_*_M_* is the sequence *x* with masked residues at all mutated position *M*. Since we only consider single amino acid substitutions in this work, *M* contains only a single position.

For a VAE, as in EVE ^19^, where the exact computation of log likelihood of a sequence is intractable, we approximate it with the Evidence Lower Bound (ELBO) used to optimize the VAE:

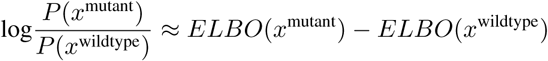

The ELBO term itself is estimated via Monte Carlo sampling, using 20k samples from the approximate posterior distribution. These approximations have been shown to provide strong results in practice ^19^. Note that this is the negative of the evolutionary index score outputted by the EVE model.

### A.7 Reliability Estimation

#### A.7.1 Alignment-based Reliability: Neff at 90% Identity

To account for uneven sampling of protein families, by both sequencing efforts and natural processes, alignment-based models reweight sequences in a multiple sequence alignment by a measure of their uniqueness. Each sequence in an alignment is assigned a weight *π_x_* = 1*/S*, where *S* is the number of sequences in that alignment within a given Hamming distance. For non-viruses this cutoff is typically set to 0.2 (80% sequence identity) ^17^. Following prior work ^2,18^, we use a cutoff of 0.01 (99% sequence identity) that works well for viral proteins, due to the relatively limited sequence diversity and an expectation that small difference in viral sequences will have comparatively large impacts on overall fitness constraints.

The effective number of non-redundant homologous sequences (Neff) is calculated using the sum of sequence weights as a measure of the sequence information present in the training alignment ^18^.

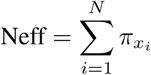

where *N* is the number of sequences in the alignment. Neff has also been expressed as the exponential of the Shannon entropy averaged over all columns of the multiple sequence alignment.

Following Hopf et al. (2017) ^18^, we divide Neff by the length of the target sequence (*L*). This ratio provides a normalized measure of sequence diversity relative to the protein length, which helps assess whether there is sufficient evolutionary information to reliably model the constraints on each position in the sequence. Alignment depth can be split into Low (Neff/L *<*1), Medium (1 *<*Neff/L *<*100), and High (Neff/L *>*100) as in ProteinGym ^17^.

As our work demonstrates the greater importance of sequences within 90% identity of the target sequence, we define an alignment-based confidence metric as the effective number of sequences within 90% identity of the target:

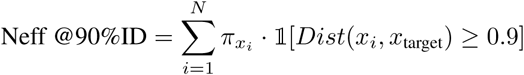

##### Alignment selection criteria

We distinguish between the metric used for confidence when comparing across viral proteins and the metric used for deciding on the optimal alignment within one viral protein (that have been made using different bitscores, databases, or other parameters). To understand the modeling choices to optimize alignment-based model performance for viruses, for each viral protein in our DMS benchmark, we generated MSAs using three reference databases: UniRef90, UniRef100, and UniRef100+BFD+MGnify, while systematically varying alignment depth at six sequence similarity thresholds during homology search (Fig. 3A). When deciding the optimal alignment for one viral protein, we demonstrate that the proportion of sequences in the alignment that are highly relevant to the target sequence (within 90% identity) is critical. We therefore choose the best alignment that has (1) Neff/L *>*1 and (2) the highest proportion of sequences within 90% identity. While this heuristic works for alignments created using a standard alignment-generating software like JackHMMER ^123^, taking this approach to the limit–maximizing the proportion of highly related sequences by selecting only sequences above the 90% identity threshold–is unlikely to be optimal. We choose 90% as the identity cutoff as the percentage in which the greatest fraction of DMSs across taxa (70%) showed positive correlation between alignment relatedness at that percentage and model performance (Fig. S11H). We also note that while alignment relevance is a better predictor of performance than fraction of viral sequences in the alignment, increasing relevance does by necessity also increase the fraction viral (Fig. S11F).

#### A.7.2 PLM Reliability: Scaled Pseudo-Perplexity

In an encoder-only PLM like ESM or SaProt, an estimation of perplexity of a sequence *x* of length *L* can be obtained by calculating the exponential of the negative pseudo-log-likelihood, or pseudo-perplexity:

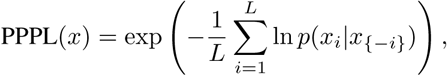

where *x*_{−*i*}_ is the set of all residues except *x_i_*^106^.

For SaProt, we calculate pseudo-perplexity only of the amino acid tokens, not structural tokens, and with the structural token unmasked when a particular amino acid token *x_i_* is masked out and predicted by the model based on all other tokens *x*_{−_*_i_*_}_. This matches the masking strategy used during training. Moreover, the structural token is left visible to reduce the model’s emphasis on predicting it, as despite inaccuracies in foldseek tokens there is still valuable context provided ^41^.

For the autoregressive PLM Tranception, we directly calculate the perplexity of a sequence *x* using the exponentiated average log-likelihood conditioned on the preceding amino acid tokens when scored from both the left and right directions:

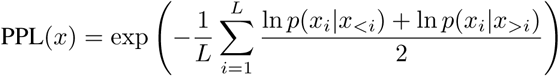

We create a PLM reliability metric calculated based on the (pseudo-)perplexity of the wildtype protein as 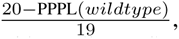 scaling the result to a value between 0 and 1, where higher scores indicate greater confidence. This scaling reflects perplexity’s interpretation as the effective number of amino acids the model is choosing between, where a perfect (pseudo-)perplexity of 1 represents complete certainty and a model completely uncertain between 20 amino acids would approach a pseudo-perplexity of 20. While we note a pseudo-perplexity of 1 represents complete certainty, in truth more than 1 amino acid should be functional at a given position and therefore there should be some uncertainty unless the model has overfit to the wildtype sequence. So, the confidence metric should rarely reach a score of 1, and a score of 0.9 would be expected and in fact desirable for high performing models. Note that negative values are technically possible with this metric, but for our purposes can be capped at 0.

We also experiment with using a “pseudo-perplexity” of the EVE Evidence Lower Bound (ELBO), a lower bound to the log-marginal likelihood of each amino acid at each position, as an alternate reliability score, though this is difficult to interpret given the overall high confidence and performance of the model.

#### A.7.3 Reliability Thresholds

We choose an optimal reliability threshold that maximizes the sensitivity and specificity (Youden’s J) for the reliability measure predicting high DMS correlation (defined as Spearman *ρ >* 0.4 between model and DMS). To calibrate these reliability estimates, we use a subset of 23 “comparable” viral DMSs. We expect correlations with DMS to be more comparable when we focus on fitness assays (measuring cell entry, replication, or infectivity), proteins with length less than 1024 (as the context size of PLMs), and studies performed after 2015 (many earlier DMSs have low replicate correlations). Given the relatively low number of DMSs to calibrate against, often a range of reliability scores will have the same sensitivity and specificity, and we choose to round down within this range. While we use Spearman *ρ >* 0.4 as a means to choose reliability thresholds here for demonstration, we expect that others will use the EVEREST benchmark as a means to define their own thresholds based on desired performance levels.

### A.8 Hybrid Models

#### A.8.1 VESPA

VESPA ^79^ combines the embeddings from ProtT5^99^ with a per-residue conservation prediction and supplements with BLOSUM substitution scores ^49^. ProtT5 uses a T5 architecture which uses an encoder and decoder and was first trained on BFD and then finetuned on UniRef50.

#### A.8.2 Tranception with MSA retrieval

Tranception ^22^ combines an autoregressive protein language model with inference-time retrieval from a MSA. We used Tranception Large (700M parameters) trained on UniRef100 as implemented in ProteinGym. For retrieval, we use the same MSA (with the bit score and database chosen via the confidence metrics) as in alignment-based methods section. The MSA is not directly trained on, but retrieval inference uses the empirical distribution of amino acids observed across sequences in the MSA retrieved set of homologous sequences calculated via pseudocounts with Laplace smoothing ^17^. Sequences are re-weighted as in Hopf et al. (2017) ^18^. The final log likelihood is a weighted average from autoregressive inference and the MSA log prior. The optimal aggregation coefficient was found to be 0.6 by grid search on a subset of DMSs.

#### A.8.3 TranceptEVE

TranceptEVE ^38^ is a hybrid method that combines Tranception ^22^ with an MSA-based EVE log prior ^19^. The same EVE log prior is used to score all sequences of interest. Unlike Tranception, the aggregation coefficients used for both the EVE log prior and the MSA log prior are dependent on the depth of the retrieved MSA for a given protein family. If the protein family of interest has no or very few homologs, the autoregressive transformer is relied on, while if the MSA is deeper, the MSA and EVE log priors are weighted higher.

#### A.8.4 SaProt-EVE

For comparison, we make a hybrid of SaProt and EVE that combines the advantages of alignment-based, family-specific models with structure-aware protein language models. We impute the EVE scores with the mean value, standard scale each of the models, shift the scores positive, and then take the geometric mean. We select the geometric mean instead of an arithmetic mean as it is less sensitive to outliers.

#### A.8.5 Evotuning SaProt

“Evotuning” is an alternative to training protein language models on new datasets, where pre-trained models are fine-tuned on a set of homologous, evolutionary sequences and generally expected to improve predictive capabilities for the targeted protein groups ^103^. We evaluated several fine-tuning strategies for the SaProt backbone on a Lassa dataset of 3102 sequences, constructed from both Foldseek and JackHMMER (optimal alignment-based model selection) searches of the Lassa virus glycoprotein complex (GPC). In all cases, the “backbone” denotes the pretrained SaProt transformer encoder and the “task head” the downstream regression head used for DMS score prediction. We considered three fine-tuning modes:

- Mode Full: full model trainable.
- Mode Unfreeze-8: backbone frozen except the last 8 layers.
- Mode Unfreeze-16: backbone frozen except the last 16 layers.

We report Spearman correlation between model predictions and experimental DMS scores. For each run we give the best single-epoch value, and where noted we also report the mean over all epochs in that run.

As an initial hyperparameter search, we performed a learning-rate sweep for Mode Unfreeze-8 with learning rate (*lr* = 10^−3^, 10^−4^, 10^−5^, 10^−6^) for 60 epochs. For each setting, we evaluated on the Lassa DMS the following models:

- Mode Unfreeze-8, *lr* = 10^−3^: average 0.19 (best 0.31)
- Mode Unfreeze-8, *lr* = 10^−4^: average 0.19 (best 0.31)
- Mode Unfreeze-8, *lr* = 10^−5^: average 0.30 (best 0.34)
- Mode Unfreeze-8, *lr* = 10^−6^: average 0.32 (best 0.34)

We therefore fixed *lr* = 10^−6^ for all subsequent SaProt experiments on Lassa-GPC. Under this setting (60 epochs), we evaluated on the Lassa DMS the following models:

- Mode Full: average 0.33 (best 0.34)
- Mode Unfreeze-16: average 0.32 (best 0.34)

To test whether span-level reconstruction improves SaProt’s representations by creating a more difficult training objective ^125^, we augmented the standard token-level masked language modeling (MLM) objective with a SpanBERT-style span prediction loss. In addition to randomly masking individual residues, we (i) sample contiguous random spans of residues and mask their internal tokens, and (ii) apply a span-boundary objective that trains the span-boundary representations to reconstruct the entire masked span. We refer to this configuration as SaProt + SpanBERT-style loss. For these runs, we fine-tuned SaProt on Lassa-GPC using Mode Full with *lr* = 10^−6^ for 60 epochs, achieving an average Spearman of 0.29 (best 0.32).

##### Dataset

We created a Lassa GPC dataset with both amino acid and foldseek tokens based on combining the JackHM-Mer selected alignment (amino acids only) used throughout this paper with a Foldseek search (amino acids and structural tokens). For the JackHMMER alignment, our goal was to assign each raw JackHMMER hit a corresponding structural sequence suitable as input to SaProt. We first assembled a pool of usable structural sequences. For each JackHMMER accession, we checked whether it matched a representative accession in the BFVD or an exact UniProt to PDB mapping, and when such a match was available we extracted the associated 3Di sequence. In parallel, we retrieved the corresponding raw amino acid sequences from the UniProt API. We then expanded this pool by running Foldseek against PDB and AF2 (UniRef50) structure databases, which yielded additional pairs of raw sequences and 3Di sequences. For all non-PDB structures, we also downloaded per residue pLDDT scores and masked 3Di tokens at positions with confidence below 0.7.

Next, we used MMseqs2 to map the original set of raw JackHMMER hits, treated as queries, to this structural sequence database. For each query, we selected the most stringent query coverage threshold in the set 70%, 60%, 50%, 40%, 30%, 20%, 10% that retained at least one match, and within the surviving hits we chose the sequence with the highest sequence identity. If a PDB hit was present whose sequence identity was at least 80% of that top hit, we instead selected the PDB sequence. Otherwise, we retained the global top identity hit. For each JackHMMER hit, we then took the chosen structural sequence and performed a global pairwise alignment to the original raw query sequence. Along this alignment, we grafted the 3Di tokens onto the query by copying tokens at aligned residue–residue positions, skipping positions aligned to gaps in the query, and inserting “#” mask tokens at query insertions or at positions that had previously been masked as low confidence because of pLDDT.

### A.9 ProteinGym

ProteinGym is a collection of benchmarks for comparing the ability of models to predict the effects of protein mutations ^17^. We used this prior benchmark to compare to non-viral DMSs and to extrapolate the results of the models tested here to the *>* 50 alignment-based, protein language model, hybrid, and inverse folding models evaluated previously (but on a more limited set of viral assays). We use ProteinGym v1.1 containing 218 deep mutational scans ^17^. Multiple sequence alignments used in ProteinGym were created for each assay by performing five search iterations of the profile HMM homology search tool JackHMMER ^123^ on the UniRef100 database of non-redundant proteins ^101^. Alignments were created using 9 bit score thresholds, from 0.1 to 0.9, and selected using structural ground truth by the highest number of significant evolutionary couplings (ECs) from EVcouplings ^126^, detailed in Notin et al. (2022) ^22^.

## B Supplementary Tables

**Supplementary Table 1:** Deep Mutational Scanning studies of 45 viral proteins across 25 papers, with fitness, stability, expression, and other assays.

| ID | Virus | Protein | Paper |
| --- | --- | --- | --- |
| *Fitness* |  |  |  |
| LASSA_GP_Carr | Lassa | GP | Carr et al. (2024)<sup>78</sup> |
| SARS2_MRPO_Flynn | SARS-CoV-2 | MRPO | Flynn et al. (2022)<sup>75</sup> |
| SARS2_BA1_SPIKE_Dadonaite | SARS-CoV-2 BA.1 | SPIKE | Dadonaite et al. (2023)<sup>50</sup> |
| SARS2_DELTA_SPIKE_Dadonaite | SARS-CoV-2 Delta | SPIKE | Dadonaite et al. (2023)<sup>50</sup> |
| DENV_POLG_Suphatrakul | Dengue virus | POLG | Suphatrakul et al. (2023)<sup>57</sup> |
| IAV_H1_HA_Doud | Influenza A virus H1N1 | H1 | Doud and Bloom (2016)<sup>63</sup> |
| IAV_H1_HA_Wu | Influenza A virus H1N1 | H1 | Wu et al. (2014)<sup>64</sup> |
| IAV_PB2_Soh | Influenza A virus H1N1 | PB2 | Soh et al. (2019)<sup>70</sup> |
| IAV_H3_HA_Lee | Influenza A virus H3N2 | H3 | Lee et al. (2018)<sup>65</sup> |
| IAV_H5_HA_Dadonaite | Influenza A virus H5N1 | H5 | Dadonaite et al. (2024)<sup>66</sup> |
| IAV_H3_NP_Doud | Influenza A virus H1N1 | NP | Doud et al. (2015)<sup>67</sup> |
| IAV_H1_NP_Doud | Influenza A virus H1N1 | NP | Doud et al. (2015)<sup>67</sup> |
| IAV_PA_Wu | Influenza A virus H1N1 | PA | Wu et al. (2015)<sup>69</sup> |
| IAV_RDRP_Li | Influenza A virus H1N1 | RDRP | Li et al. (2023)<sup>71</sup> |
| NIPAH_F_Larsen | Nipah Henipavirus | F | Larsen et al. (2025)<sup>77</sup> |
| AAV2_CAPSD_Sinai | AAV2 | CAPSD | Sinai et al. (2021)<sup>54</sup> |
| CVB3_2A_Alvarez | Coxsackievirus B3 | 2A | Álvarez-Rodríguez et al. (2024)<sup>56</sup> |
| CVB3_2B_Alvarez | Coxsackievirus B3 | 2B | Álvarez-Rodríguez et al. (2024)<sup>56</sup> |
| CVB3_2C_Alvarez | Coxsackievirus B3 | 2C | Álvarez-Rodríguez et al. (2024)<sup>56</sup> |
| CVB3_3A_Alvarez | Coxsackievirus B3 | 3A | Álvarez-Rodríguez et al. (2024)<sup>56</sup> |
| CVB3_3B_Alvarez | Coxsackievirus B3 | 3B | Álvarez-Rodríguez et al. (2024)<sup>56</sup> |
| CVB3_3C_Alvarez | Coxsackievirus B3 | 3C | Álvarez-Rodríguez et al. (2024)<sup>56</sup> |
| CVB3_3D_Alvarez | Coxsackievirus B3 | 3D | Álvarez-Rodríguez et al. (2024)<sup>56</sup> |
| CVB3_VP1_Alvarez | Coxsackievirus B3 | VP1 | Álvarez-Rodríguez et al. (2024)<sup>56</sup> |
| CVB3_VP3_Alvarez | Coxsackievirus B3 | VP3 | Álvarez-Rodríguez et al. (2024)<sup>56</sup> |
| EV_CAPSD_Bakhache | Enterovirus A | CAPSD | Bakhache et al. (2024)<sup>59</sup> |
| EV_REP_Bakhache | Enterovirus A | REP | Bakhache et al. (2024)<sup>59</sup> |
| CVB3_POLG_Mattenberger | Coxsackievirus B3 | POLG | Mattenberger et al. (2021)<sup>55</sup> |
| HIV1_BF520_ENV_Haddox | HIV-1 BF520 | ENV | Haddox et al. (2018)<sup>60</sup> |
| HIV1_HV1B9_ENV_DuenasDecamp | HIV-1 HV1B9 | ENV | Duenas-Decamp et al. (2016)<sup>61</sup> |
| HIV1_BG505_ENV_Haddox | HIV-1 BG505 | ENV | Haddox et al. (2018)<sup>60</sup> |
| *Stability* |  |  |  |
| PESV_POLG_Tsuboyama | Porcine enteric sapovirus | POLG | Tsuboyama et al. (2023)<sup>58</sup> |
| BPP22_COAT_Tsuboyama | Salmonella phage P22 | COAT | Tsuboyama et al. (2023)<sup>58</sup> |
| BPT7_RPOL_Tsuboyama | Enterobacteria phage T7 | RPOL | Tsuboyama et al. (2023)<sup>58</sup> |
| LAMBDA_HCP_Tsuboyama | Enterobacteria phage lambda | HCP | Tsuboyama et al. (2023)<sup>58</sup> |
| BP434_RPC1_Tsuboyama | Enterobacteria phage 434 | RPC1 | Tsuboyama et al. (2023)<sup>58</sup> |
| *Expression* |  |  |  |
| SARS2_PRD0038_RBD_Starr | Bat coronavirus PRD0038 | RBD | Starr (2024)<sup>73</sup> |
| RmYN02_RBD_Starr | Bat coronavirus RmYN02 | RBD | Starr (2024)<sup>73</sup> |
| RsYN04_RBD_Starr | Bat coronavirus RsYN04 | RBD | Starr (2024)<sup>73</sup> |
| SARS2_XBB15_RBD_Taylor | SARS-CoV-2 XBB.1.5 | RBD | Taylor and Starr (2023)<sup>74</sup> |
| SARS2_RBD_Starr_expression | SARS-CoV-2 | RBD | Starr et al. (2020)<sup>4</sup> |

| **ID** | **Virus** | **Protein** | **Paper** |
| --- | --- | --- | --- |
| *Binding* |  |  |  |
| SARS2_RBD_Starr.binding | SARS-CoV-2 | RBD | Starr et al. (2020) <sup>4</sup> |
| IAV_NA_Jiang | Influenza A virus H3N2 | NA | Jiang et al. (2016) <sup>68</sup> |
| *Activity* |  |  |  |
| SARS2_PLPRO_Wu.activity | SARS-CoV-2 | PLPRO | Wu et al. (2024) <sup>76</sup> |
| *Abundance* |  |  |  |
| SARS2_PLPRO_Wu.abundance | SARS-CoV-2 | PLPRO | Wu et al. (2024) <sup>76</sup> |

**Supplementary Table 2:** Model performance previously available in ProteinGym ^17^. Note this analysis only covers half of the now curated 45 viral datasets and does not include SaProt-PDB, but can be used to contextualize our findings in terms of 44 models.

| Model Type | Model | Avg Spearman | Viral Avg Spearman |
| --- | --- | --- | --- |
| Alignment-based model | EVE (ensemble) | 0.439 | 0.428 |
|  | EVE (single) | 0.433 | 0.424 |
|  | DeepSequence (ensemble) | 0.419 | 0.344 |
|  | DeepSequence (single) | 0.407 | 0.323 |
|  | Potts model | 0.395 | 0.388 |
|  | Site-Independent | 0.359 | 0.383 |
| Hybrid - Alignment & PLM | TranceptEVE L | 0.456 | 0.453 |
|  | TranceptEVE M | 0.455 | 0.441 |
|  | TranceptEVE S | 0.452 | 0.433 |
|  | Tranception L | 0.434 | 0.432 |
|  | MSA Transformer (ensemble) | 0.434 | 0.414 |
|  | Tranception M | 0.427 | 0.415 |
|  | MSA Transformer (single) | 0.421 | 0.390 |
|  | Tranception S | 0.418 | 0.405 |
|  | Tranception M no retrieval | 0.348 | 0.349 |
|  | Unirep evotuned | 0.347 | 0.349 |
| Hybrid - Structure & PLM | SaProt-AF2 (650M) | 0.457 | 0.300 |
|  | ProtSSN (ensemble) | 0.449 | 0.356 |
|  | ProtSSN (k=20 h=1280) | 0.442 | 0.347 |
|  | ProtSSN (k=20 h=512) | 0.441 | 0.359 |
| Hybrid - other | VESPA | 0.436 | 0.432 |
|  | VESPAI | 0.394 | 0.392 |
| Protein language model | ProGen2 XL | 0.391 | 0.391 |
|  | ProGen2 L | 0.380 | 0.333 |
|  | ProGen2 M | 0.379 | 0.342 |
|  | ProGen2 Base | 0.378 | 0.328 |
|  | Tranception L no retrieval | 0.374 | 0.395 |
|  | ESM-1v (ensemble) | 0.407 | 0.279 |
|  | ESM-1v (single) | 0.374 | 0.258 |
|  | CARP (640M) | 0.368 | 0.273 |
|  | RITA XL | 0.372 | 0.402 |
|  | RITA L | 0.365 | 0.391 |
|  | RITA M | 0.350 | 0.385 |
|  | ProGen2 S | 0.336 | 0.285 |
|  | CARP (76M) | 0.328 | 0.150 |
|  | ESM-2 (3B) | 0.406 | 0.274 |
|  | ESM-2 (15B) | 0.401 | 0.313 |
|  | ESM-2 (150M) | 0.387 | 0.137 |
|  | ESM-2 (650M) | 0.414 | 0.238 |
|  | ESM-2 (35M) | 0.321 | 0.102 |
| Inverse folding model | ESM-IF1 | 0.422 | 0.374 |
|  | MIF | 0.383 | 0.359 |
|  | ProteinMPNN | 0.258 | 0.248 |
| Tree-based model | GEMME | 0.455 | 0.469 |

**Supplementary Table 3:** Single-sequence Protein Language Model performance of models in ProteinGym ^17^ with model details collected from model references. No hybrid models or retrieval included. Training objectives are autoregressive (AR) and masked language model (MLM); all architectures are transformer-based besides CARP which is CNN-based and Unirep which is an RNN. Performance recorded as average Spearman across ProteinGym DMSs–this analysis only covers half of the now curated 45 viral datasets, but can be used to contextualize our findings. Data size in approximate number of sequences (note different databases have different sequence length distributions ^99^). Date cutoffs and dataset splits influence training data size even when same database is used. Some model details were estimated from published papers when not explicitly stated by the authors. Ens.: Ensemble.

| Model | Parameters | Data Size | Objective | Cutoff | Context | All Taxa ($\rho$) | Virus ($\rho$) | Reference |
| --- | --- | --- | --- | --- | --- | --- | --- | --- |
| *Uniref100* |  |  |  |  |  |  |  |  |
| RITA XL | 1.2B | 280M | AR | 2022 | 1024 | 0.372 | 0.402 | Hesslow et al. (2022)<sup>23</sup> |
| Tranception L | 700M | 250M | AR | 2022 | 1024 | 0.374 | 0.395 | Notin et al. (2022)<sup>22</sup> |
| RITA L | 680M | 280M | AR | 2022 | 1024 | 0.365 | 0.391 | Hesslow et al. (2022)<sup>23</sup> |
| RITA M | 300M | 280M | AR | 2022 | 1024 | 0.350 | 0.385 | Hesslow et al. (2022)<sup>23</sup> |
| RITA S | 85M | 280M | AR | 2022 | 1024 | 0.304 | 0.358 | Hesslow et al. (2022)<sup>23</sup> |
| Tranception M | 300M | 250M | AR | 2022 | 1024 | 0.348 | 0.349 | Notin et al. (2022)<sup>22</sup> |
| Tranception S | 85M | 250M | AR | 2022 | 1024 | 0.303 | 0.315 | Notin et al. (2022)<sup>22</sup> |
| *Uniref90+BFD30* |  |  |  |  |  |  |  |  |
| ProGen2 XL | 6.4B | 230M | AR | 2022 | 1024 | 0.391 | 0.391 | Nijkamp et al. (2023)<sup>24</sup> |
| ProGen2 L | 2.7B | 230M | AR | 2022 | 1024 | 0.380 | 0.333 | Nijkamp et al. (2023)<sup>24</sup> |
| ProGen2 M | 764M | 230M | AR | 2022 | 1024 | 0.379 | 0.342 | Nijkamp et al. (2023)<sup>24</sup> |
| ProGen2 Base | 764M | 230M | AR | 2022 | 2048 | 0.378 | 0.328 | Nijkamp et al. (2023)<sup>24</sup> |
| ProGen2 S | 151M | 230M | AR | 2022 | 1024 | 0.336 | 0.285 | Nijkamp et al. (2023)<sup>24</sup> |
| *Uniref90* |  |  |  |  |  |  |  |  |
| ESM-2 (650M) | 650M | 65M | MLM | 2022 | 1024 | 0.414 | 0.238 | Lin et al. (2023)<sup>25</sup> |
| ESM-1v (ens.) | 650M | 98M | MLM | 2020 | 1024 | 0.407 | 0.279 | Meier et al. (2021)<sup>21</sup> |
| ESM-2 (3B) | 3B | 65M | MLM | 2022 | 1024 | 0.406 | 0.274 | Lin et al. (2023)<sup>25</sup> |
| ESM-2 (15B) | 15B | 65M | MLM | 2022 | 1024 | 0.401 | 0.313 | Lin et al. (2023)<sup>25</sup> |
| ESM-2 (150M) | 150M | 65M | MLM | 2022 | 1024 | 0.387 | 0.137 | Lin et al. (2023)<sup>25</sup> |
| ESM-1v | 650M | 98M | MLM | 2020 | 1024 | 0.374 | 0.245 | Meier et al. (2021)<sup>21</sup> |
| ESM-2 (35M) | 35M | 65M | MLM | 2022 | 1024 | 0.321 | 0.102 | Lin et al. (2023)<sup>25</sup> |
| ESM-2 (8M) | 8M | 65M | MLM | 2022 | 1024 | 0.226 | 0.078 | Lin et al. (2023)<sup>25</sup> |
| *Uniref50* |  |  |  |  |  |  |  |  |
| ESM-1b | 650M | 27M | MLM | 2018 | 1024 | 0.394 | 0.241 | Rives et al. (2021)<sup>127</sup> |
| CARP (640M) | 640M | 40M | MLM | 2023 | 4096 | 0.368 | 0.273 | Yang et al. (2024)<sup>26</sup> |
| CARP (76M) | 76M | 40M | MLM | 2023 | 4096 | 0.328 | 0.150 | Yang et al. (2024)<sup>26</sup> |
| CARP (38M) | 38M | 40M | MLM | 2023 | 4096 | 0.279 | 0.125 | Yang et al. (2024)<sup>26</sup> |
| ProtGPT2 | 738M | 40M | AR | 2022 | 1024 | 0.188 | 0.141 | Ferruz et al. (2022)<sup>128</sup> |
| Unirep | 18.2M | 24M | AR | 2018 | 280 | 0.190 | 0.057 | Alley et al. (2019)<sup>129</sup> |
| CARP (600K) | 600K | 40M | MLM | 2023 | 4096 | 0.106 | 0.042 | Yang et al. (2024)<sup>26</sup> |

**Supplementary Table 4:** Number of successful Potts models given the equal memory constraints at different bitscores across UniRef90, UniRef100, and UniRef100+MGnify+BFD datasets. Failed models with high numbers of sequences would mostly be excluded by selected Neff cutoff, as only a subset of bit scores/databases failed per viral DMS.

| Bitscore | UniRef90 | UniRef100 | UniRef100+MGnify+BFD |
| --- | --- | --- | --- |
| 0.5 | 57 | 57 | 48 |
| 0.4 | 57 | 56 | 37 |
| 0.3 | 55 | 51 | 28 |
| 0.05 | 54 | 51 | 21 |
| 0.04 | 53 | 45 | 12 |
| 0.03 | 44 | 37 | 10 |

**Supplementary Table 5:**
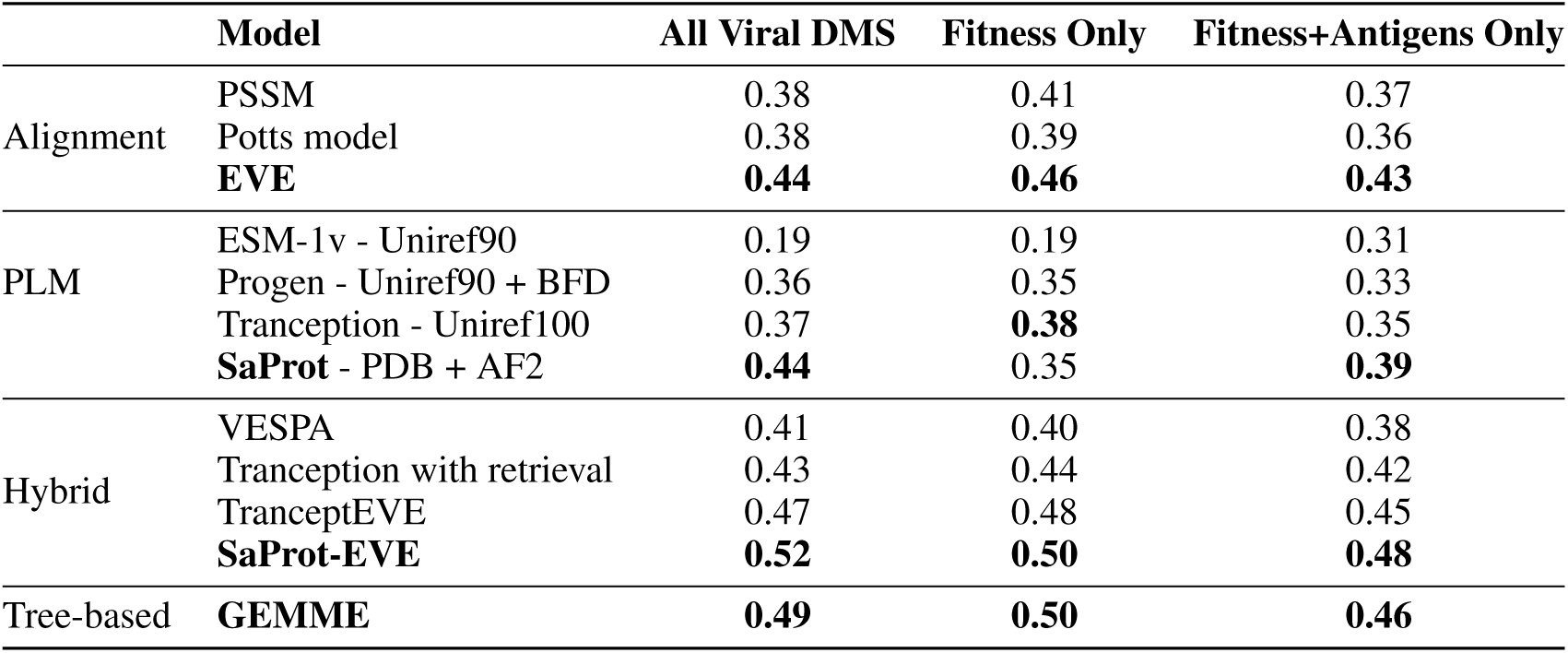
Model Performance (average Spearman *ρ*) on EVEREST DMS Benchmark with 45 viral DMSs, or on the subset of scans with an encompassing fitness assay measuring infectivity, replication, or cell entry, or the subset of fitness assays that are antigens.

**Supplementary Table 6:** MSA (Neff @ 90% identity) and PLM (scaled pseudo-perplexity, SaProt) reliability scores for 44 antigens of 40 WHO priority and prototype RNA viruses. RBP: Receptor Binding Protein, AP: Accompanying Protein.

| ID | Family | Structural Class | MSA confidence | PLM confidence |
| --- | --- | --- | --- | --- |
| Junin_GPC | Arena | Class I Fusion | 5 | 0.61 |
| Lassa_GPC | Arena | Class I Fusion | 259 | 0.55 |
| Lujo_GPC | Arena | Class I Fusion | 1 | 0.60 |
| MERS_Spike | Corona | Class I Fusion | 8 | 0.67 |
| SARS2_Spike | Corona | Class I Fusion | 157 | 0.67 |
| Ebola_GP | Filo | Class I Fusion | 85 | 0.70 |
| Marburg_GP | Filo | Class I Fusion | 15 | 0.57 |
| SEBOV_GP | Filo | Class I Fusion | 3 | 0.68 |
| DENV_E | Flavi | Class II Fusion | 107 | 0.90 |
| TBE_E | Flavi | Class II Fusion | 98 | 0.81 |
| WNV_E | Flavi | Class II Fusion | 11 | 0.82 |
| YFV_E | Flavi | Class II Fusion | 35 | 0.83 |
| Zika_E | Flavi | Class II Fusion | 50 | 0.90 |
| HTNV_Gc | Hanta | Class II Fusion | 13 | 0.69 |
| HTNV_Gn | Hanta | Class II AP | 12 | 0.53 |
| SNV_Gc | Hanta | Class II Fusion | 13 | 0.71 |
| HEV3_orf2 | Hepe | Capsid | 128 | 0.67 |
| IAV_H1 | Orthomyxo | Class I Fusion | 556 | 0.90 |
| IAV_H10 | Orthomyxo | Class I Fusion | 164 | 0.82 |
| IAV_H2 | Orthomyxo | Class I Fusion | 184 | 0.91 |
| IAV_H3 | Orthomyxo | Class I Fusion | 349 | 0.88 |
| IAV_H5 | Orthomyxo | Class I Fusion | 638 | 0.90 |
| IAV_H6 | Orthomyxo | Class I Fusion | 226 | 0.88 |
| IAV_H7 | Orthomyxo | Class I Fusion | 238 | 0.87 |
| NiV_F | Paramyxo | Class I Fusion | 2 | 0.69 |
| NiV_G | Paramyxo | Non-fusion RBP | 9 | 0.73 |
| OROV_G1 | Peribunya | Class II Fusion | 8 | 0.62 |
| OROV_G2 | Peribunya | Class II AP | 8 | 0.60 |
| RVF_Gn | Phenui | Class II AP | 29 | 0.57 |
| SFTS_Gc | Phenui | Class II Fusion | 9 | 0.73 |
| SFTS_Gn | Phenui | Class II AP | 35 | 0.58 |
| hPBV_capsid | Picobirna | Capsid | 2 | 0.51 |
| hPBV_capsid | Picobirna | Capsid | 2 | 0.51 |
| EV-D68_VP1 | Picorna | Capsid | 85 | 0.79 |
| EV71_VP1 | Picorna | Capsid | 269 | 0.85 |
| polio_capsid | Picorna | Capsid | 13 | 0.84 |
| hMPV_F | Pneumo | Class I Fusion | 39 | 0.74 |
| hMPV_G | Pneumo | Non-fusion RBP | 2 | 0.48 |
| HIV1_Env | Retro | Class I Fusion | 20 | 0.87 |
| VSV_GP | Rhabdo | Class III Fusion | 17 | 0.61 |
| CHIKV_E1 | Toga | Class II Fusion | 32 | 0.84 |
| CHIKV_E2 | Toga | Class II AP | 49 | 0.77 |
| VEE_E1 | Toga | Class II Fusion | 11 | 0.78 |
| VEE_E2 | Toga | Class II AP | 15 | 0.74 |

## C Supplementary Figures

**Supplementary Figure 1:**
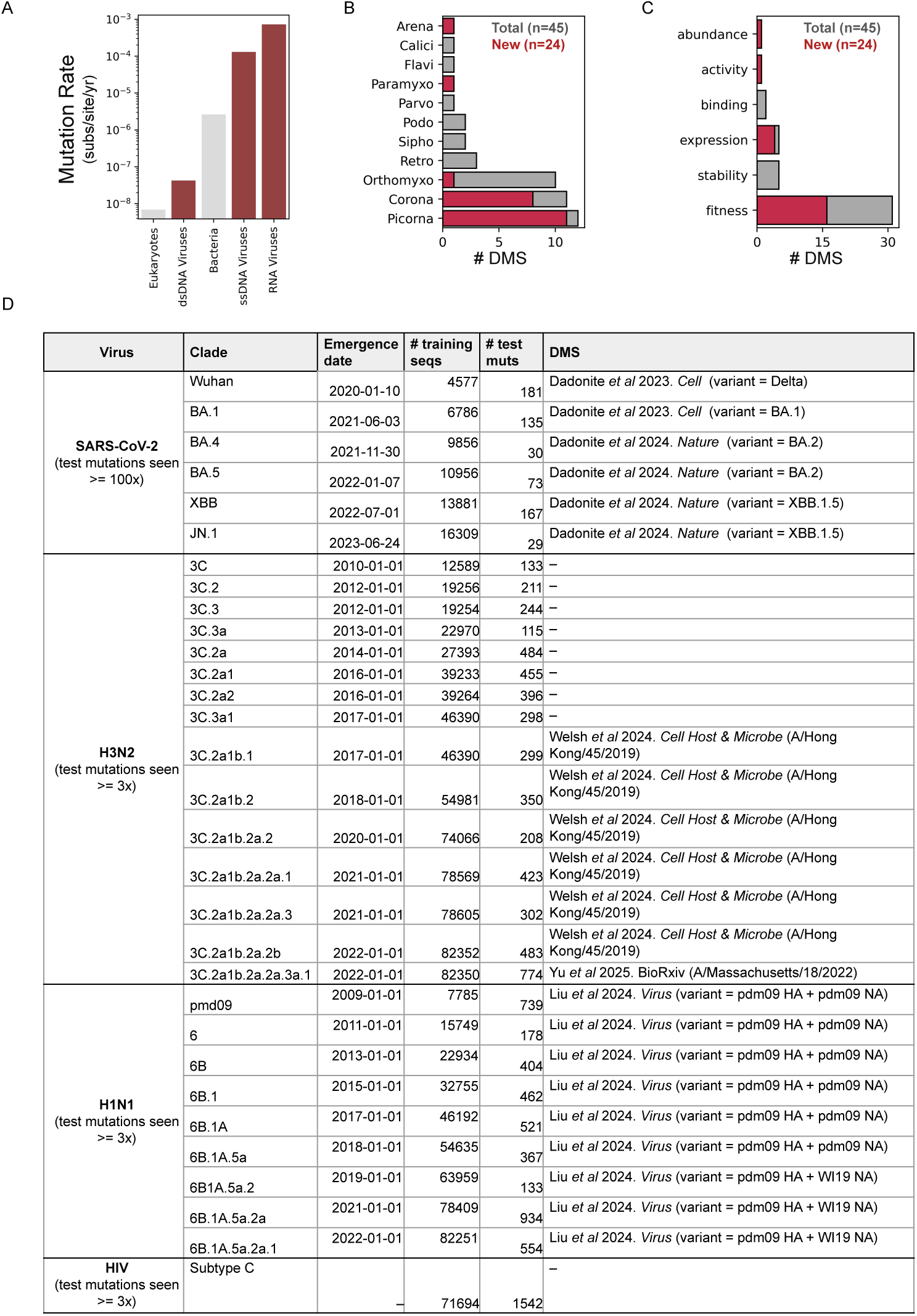
Overview of EVEREST virus-specific benchmarking dataset. (A) Viruses, especially RNA viruses, require curated benchmarking because of unique features including have higher mutation rates than other organisms. Geometric average mutation rates (substitutions per site per year) from Linz et al. ^47^. (B-C) EVEREST DMS benchmark comprised of 45 experimental scans, approximately doubling the number of viral assays included in prior benchmarks ^17^. EVEREST covers 11 viral families (B) and diverse assay phenotypes (C). (D) EVEREST real-work forecasting benchmark, includes 31 clades from 4 virus: SARS-CoV-2, H3N2, H1N1, and HIV, and compares the forecasting ability of fitness DMSs (matched to the starting clade when possible) ^5,50–53^.

**Supplementary Figure 2:**
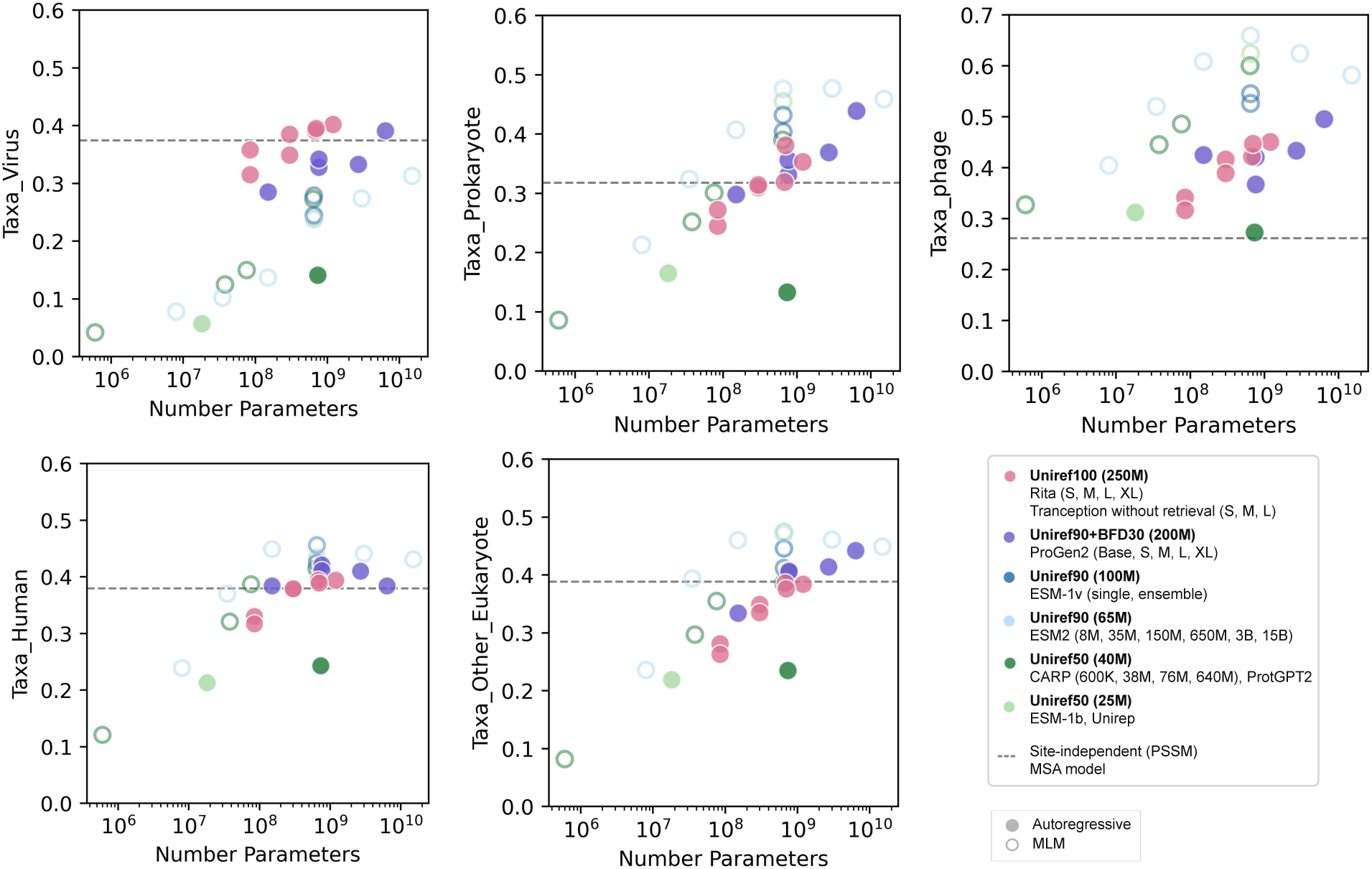
Relationship between protein language model size and training dataset and model performance (average Spearman) across different taxa. Scaling number of PLM parameters leads to higher average Spearman correlation with viral proteins (top left). The best PLMs for viruses are autoregressive with larger training datasets, while masked language models generally have better performance on other taxa. Phages (also all stability assays) separated out from eukaryotic-infecting viruses due to differences in performance. Most PLMs outperform site-independent alignment-based model (dotted line) for all taxa except viruses.

**Supplementary Figure 3:**
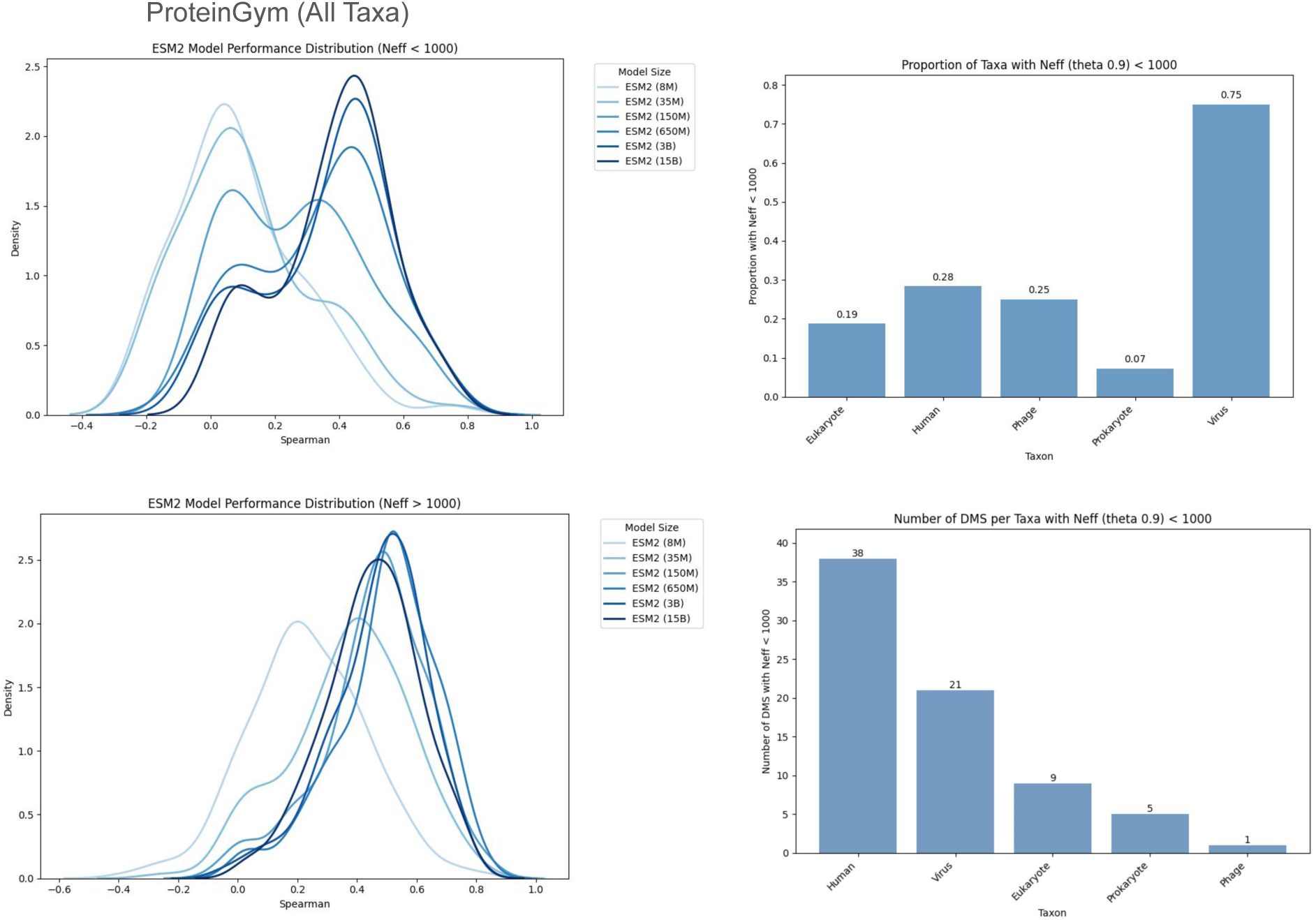
The masked protein language model ESM2 benefits from increased number of parameters especially when there is a low number of related sequences in the PLM training data (calculated as number of effective sequences in a JackHMMER alignment of Uniref100 with clustering at theta 90%, since ESM2 is trained on Uniref90) when looking DMSs from all taxa in ProteinGym. This impacts the majority of viruses as 75% of the viruses in ProteinGym have Neff less than 1000, while between 7% and 28% of other taxa have such low effective number of sequences post-clustering (Neff; though the low Neff group has a substantial number of non-viruses, as in total there are more than 5 times the number of non-viral DMS as viral; bottom right). When Neff is greater than 1000, once the number of parameters is at least 150M there is little difference in performance as model size increases.

**Supplementary Figure 4:**
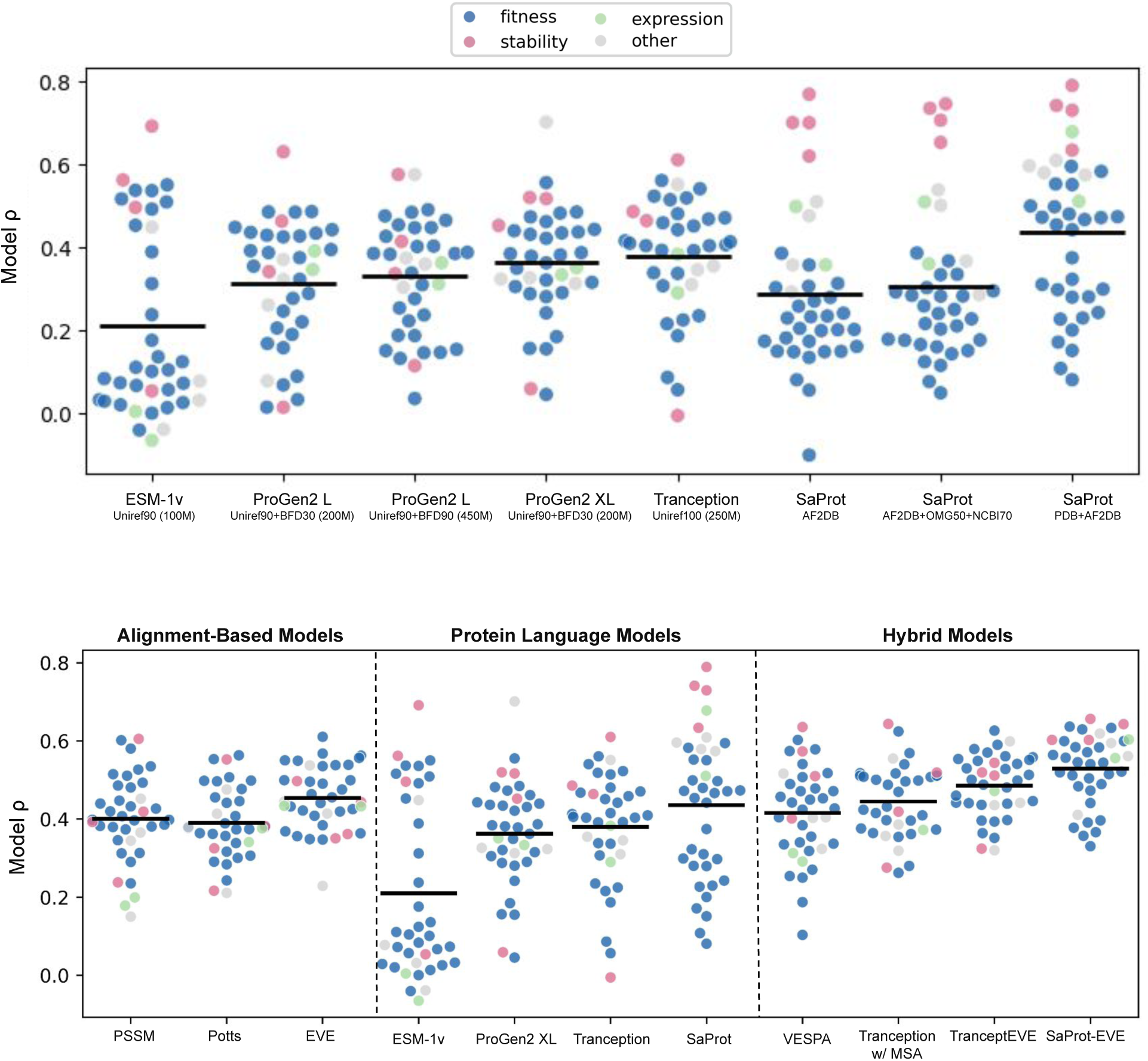
Performance of alignment-based, protein language models, and hybrids on EVEREST DMS benchmark.

**Supplementary Figure 5:**
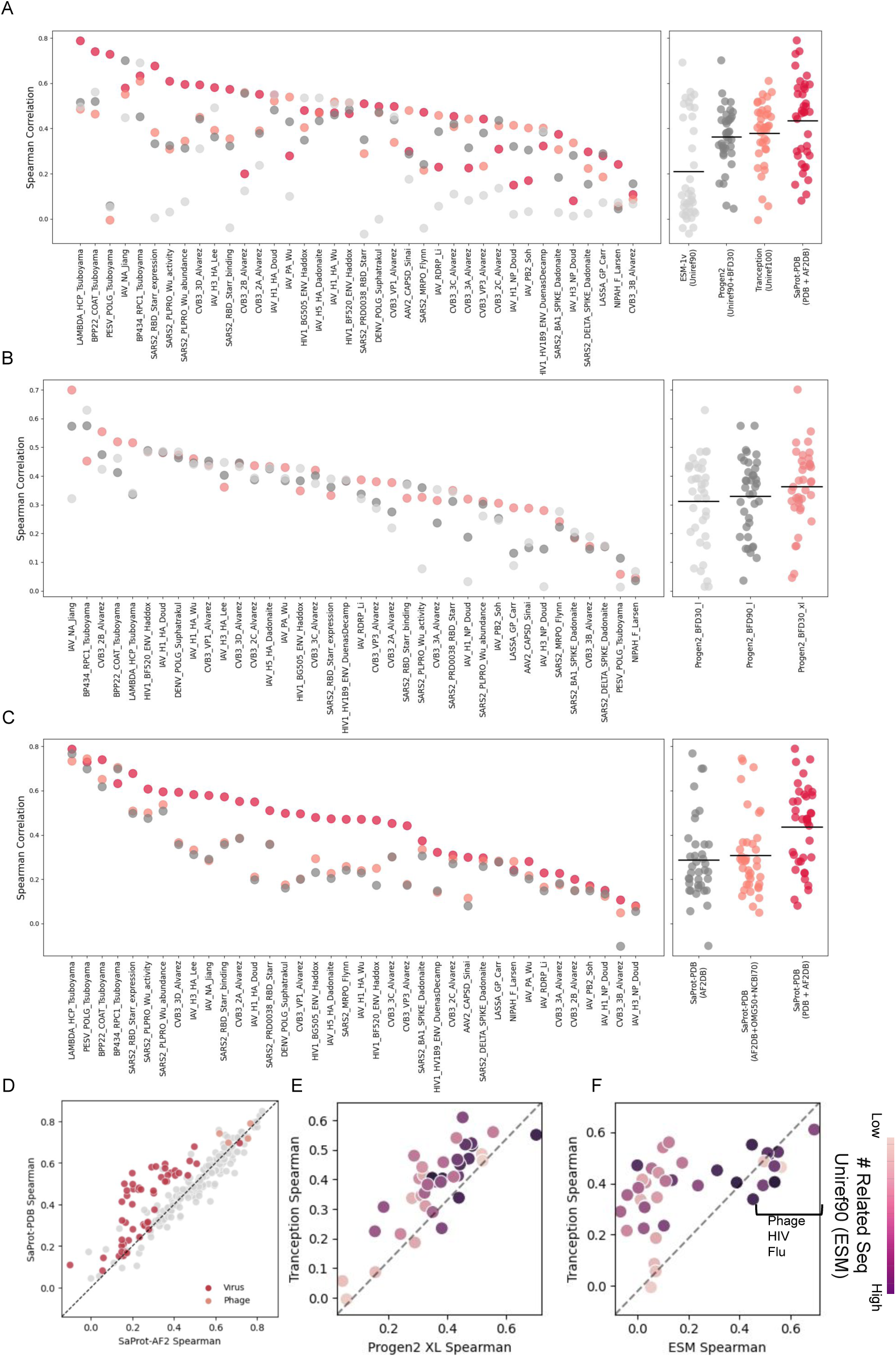
Breakdown of protein language model performance on each protein in EVEREST DMS benchmark. (A) Performance of models shown in main: ESM-1v, ProGen2, Tranception and SaProt. (B) Performance of different ProGen2 models with different model size and training datasets. (C) Performance of different SaProt models with different training datasets. (D) Comparison of the Spearman rank correlation of SaProt-PDB to SaProt-AF2 for all DMSs from ProteinGym. SaProt-PDB outperforms SaProt-AF2 for most eukaryotic viral proteins, but not for phages. (E-F) Differences in performance between Tranception and ProGen2 (E) or ESM-1v (F) colors by number of related sequences in Uniref90 training dataset.

**Supplementary Figure 6:**
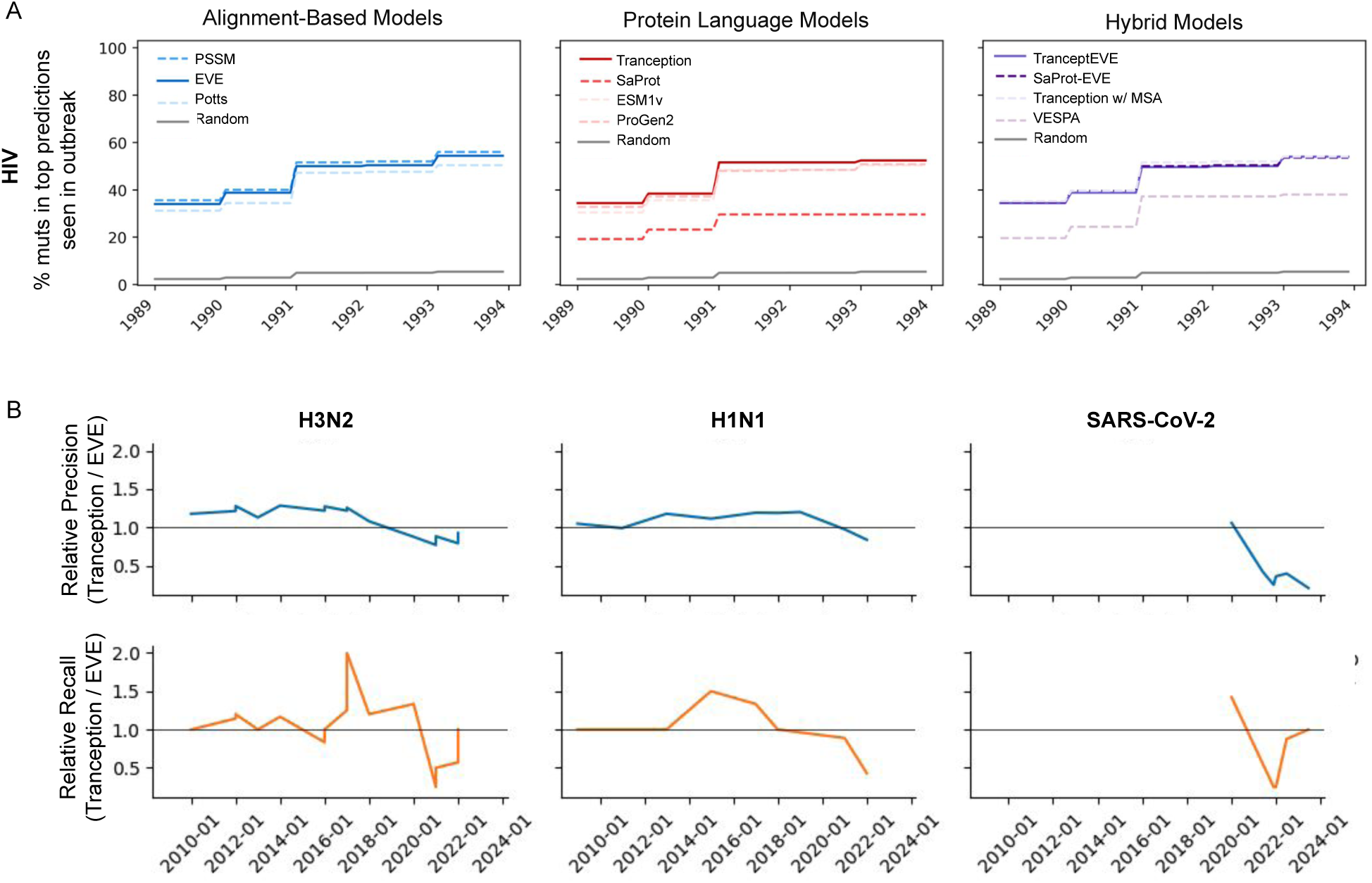
Model performance on EVEREST real-world viral evolution forecasting benchmark. (A) Time-course analyses shown for HIV clade. Model predictions correspond to top 250 ranked mutations. Highly frequent mutations are those observed *>*1k times for influenza and *>*10k times for SARS-CoV-2 and HIV. (B) Relative performance of of Tranception versus EVE across time. Top panel is relative performance of model-predicted mutations subsequently occurring during evolution. Bottom panel is relative performance on model recalling frequent mutations.

**Supplementary Figure 7:**
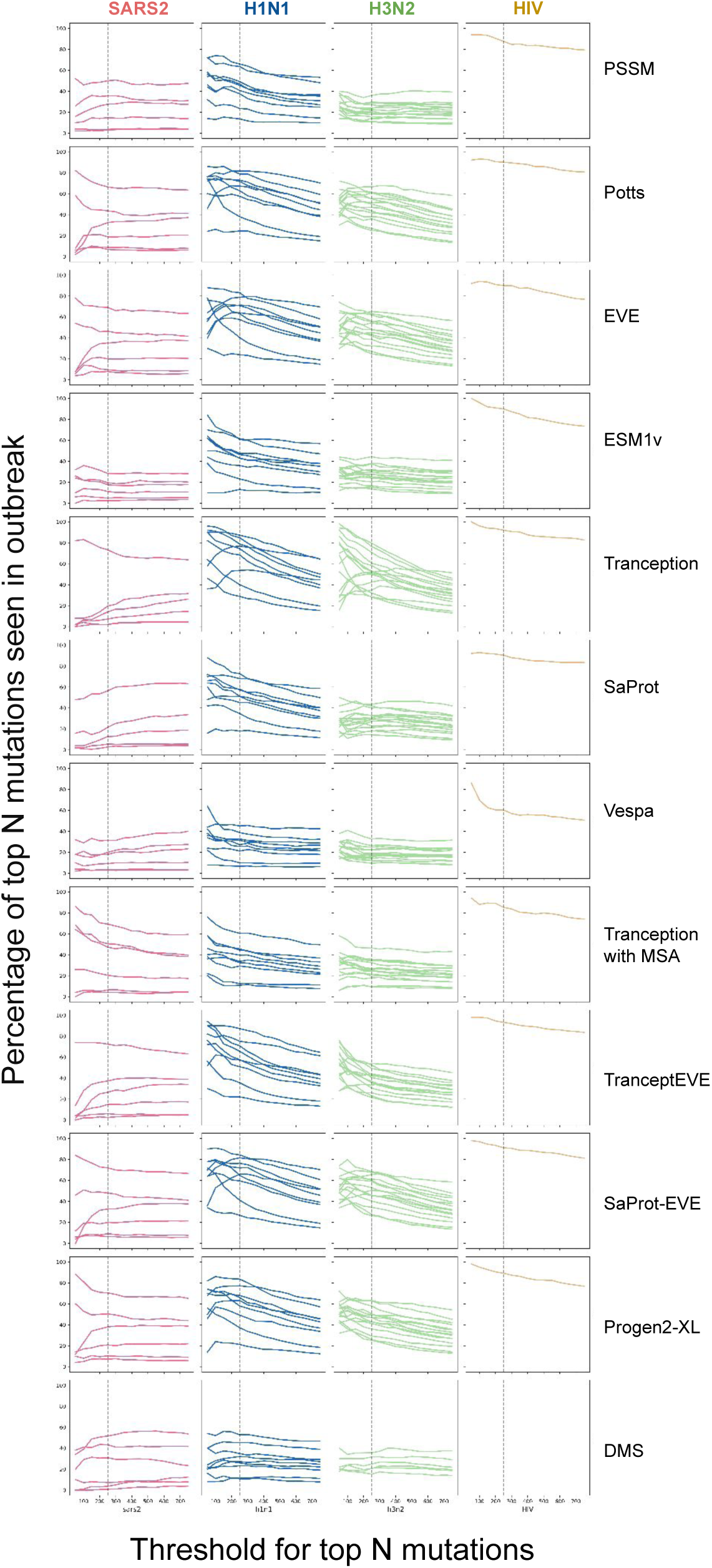
Model performance at different top prediction cutoffs. Performance measured as the percentage of mutations seen during evolution. Each column is a virus with 6 SARS-CoV-2 clades, 9 H1N1 clades, 15 H3N2 clades and 1 HIV clade. Highly frequent mutations are those observed *>*1k times for influenza and *>*10k times for SARS-CoV-2 and HIV.

**Supplementary Figure 8:**
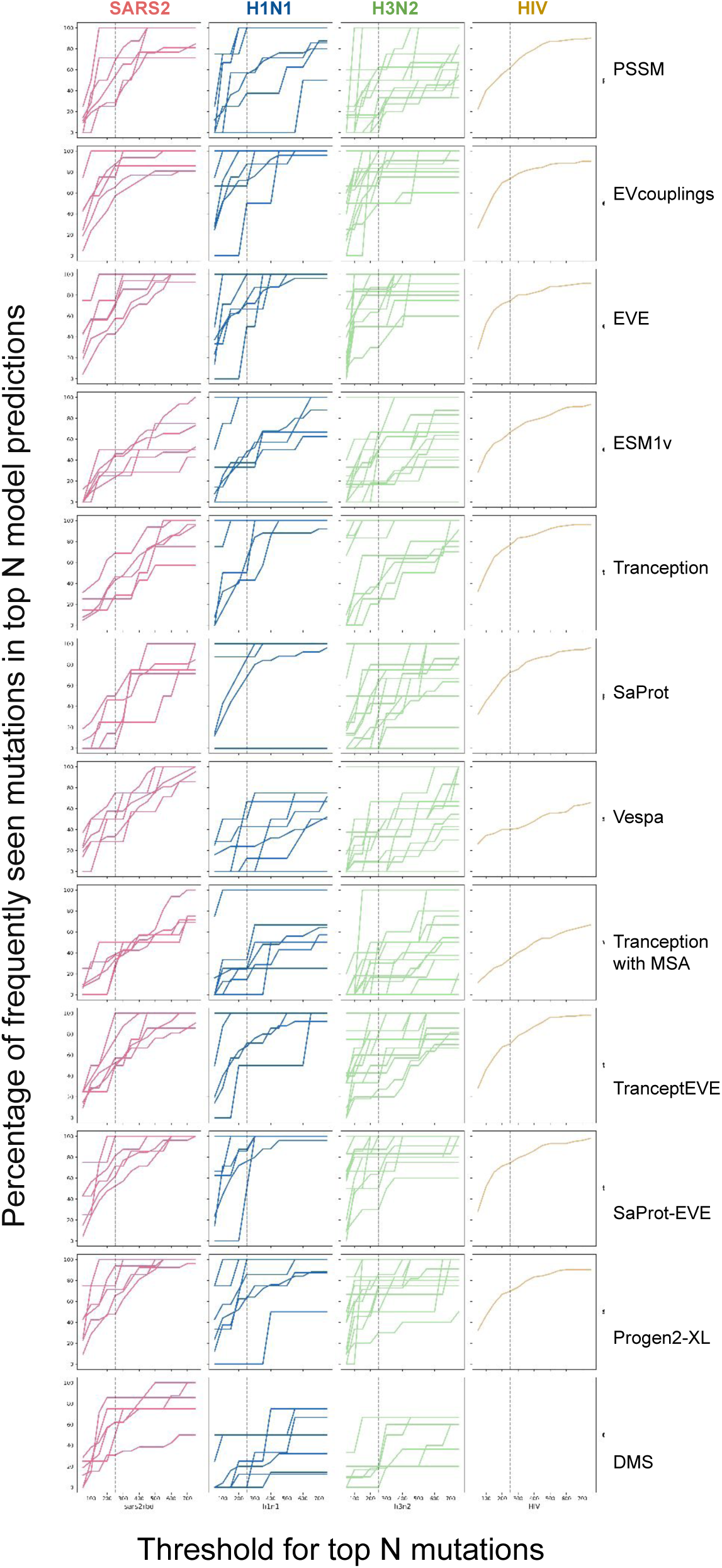
Model performance at different top prediction cutoffs. Performance measured as percentage of highly frequent mutations captured by model predictions. Each column is a virus with 6 SARS-CoV-2 clades, 9 H1N1 clades, 15 H3N2 clades and 1 HIV clade. Highly frequent mutations are those observed *>*1k times for influenza and *>*10k times for SARS-CoV-2 and HIV.

**Supplementary Figure 9:**
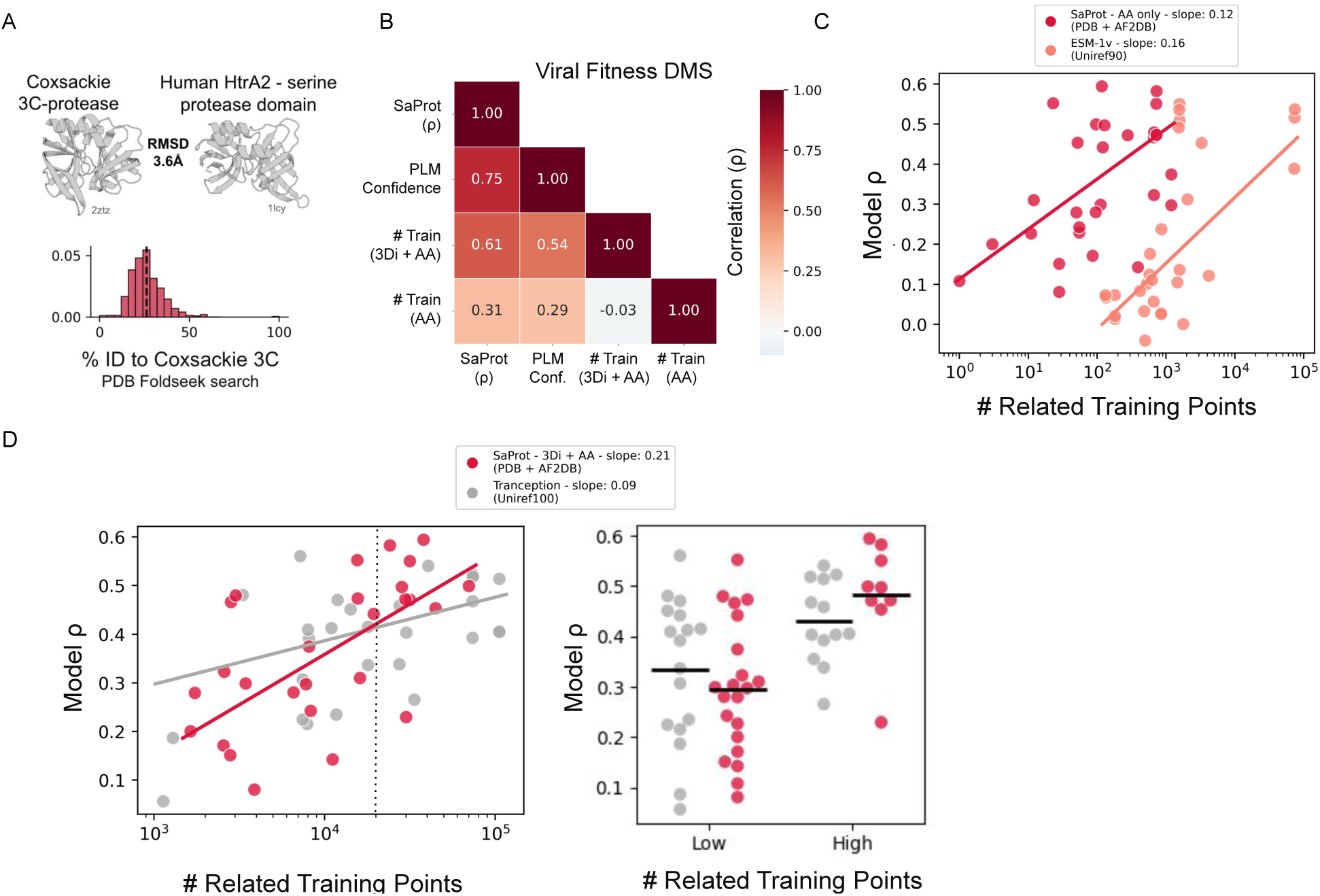
(A) By incorporating structural information, many more homologous proteins can be retrieved than by sequence alone when using comparable approaches—between 10^3^ − 10^5^ using Foldseek versus fewer than 10^3^ using MMseqs2. For instance, a Foldseek search using coxsackievirus 3C protease retrieved thousands of structurally related proteins, most with less than 50% sequence identity. Among them was human HtrA2, a serine protease which shares structural similarity with cysteine proteases (such as the coxsackievirus 3C protease) across the tree of life ^130^ and membership in the same CATH and SCOP superfamilies ^131,132^ (Fig. S9). Whether this is an example of true homology (divergence from a common ancestor) or convergent evolution on a common biochemistry is a matter of debate ^133^. Coxsackievirus 3C protease structure (2ztz chain A) and Human HtrA2 serine protease domain (1lcy, selected residues 1-210^134^) share a common fold with RMSD of 3.6Å. RMSD identified by CE algorithm for alignment, defined over 160 residues, robust for structures with low sequence similarity. Pairwise sequence identity of all sequences from the Foldseek (3Di + AA, n=2164) search of Coxsackievirus 3C in the PDB, dotted line shown for HtrA2. While such remote homologs are retrievable with more robust sequence homology methods like JackHMMER, their great distance in sequence space may prevent sequence-only PLMs from learning their relevance, and could benefit from structure tokens to increase this signal. These remote structural analogs therefore likely contribute to SaProt’s improved generalization across divergent protein families given sufficient structure information. (B) Correlations across viral fitness DMS benchmark between SaProt performance (Spearman correlation with DMS), SaProt pseudo-perplexity-based reliability score, number of related sequences in training data identified via sequence homology search, and number or related sequences in training data identified via sequence and structural homology search. (C) PLM performance corresponds to number of relevant sequences in training datasets (via mmseqs search of amino acids (AA) only). SaProt (trained on PDB + AF2DB) learns from much fewer related sequences (by amino acid identity) than ESM-1v (trained on UniRef90) to achieve greater correlation with DMS, despite using an ESM architecture. (D) PLM performance heavily corresponds to number of relevant sequences and structures in training datasets (via Foldseek search using amino acids and 3Di tokens) (left). SaProt may have a performance boost or be more consistent when there are high numbers of related training points (though this is less common to have high numbers of relevant training points in the PDB+AF2DB). Swarmplot (right) shown when splitting number of relevant sequences for each appropriate training dataset based on whether greater than 20,000 (dotted line on left). Slopes calculated with robust linear model estimation using RANSAC. However, the extent to which errors arise in predicted viral structures resulting from low sequence diversity ^105^ remains to be investigated and may impact downstream performance, even when only local structural information is encoded. Moreover, it is also possible that learning from the PDB has been optimized such that its few data points are exceptionally useful for mutation effect prediction as more important sequences are generally selected for structural characterization.

**Supplementary Figure 10:**
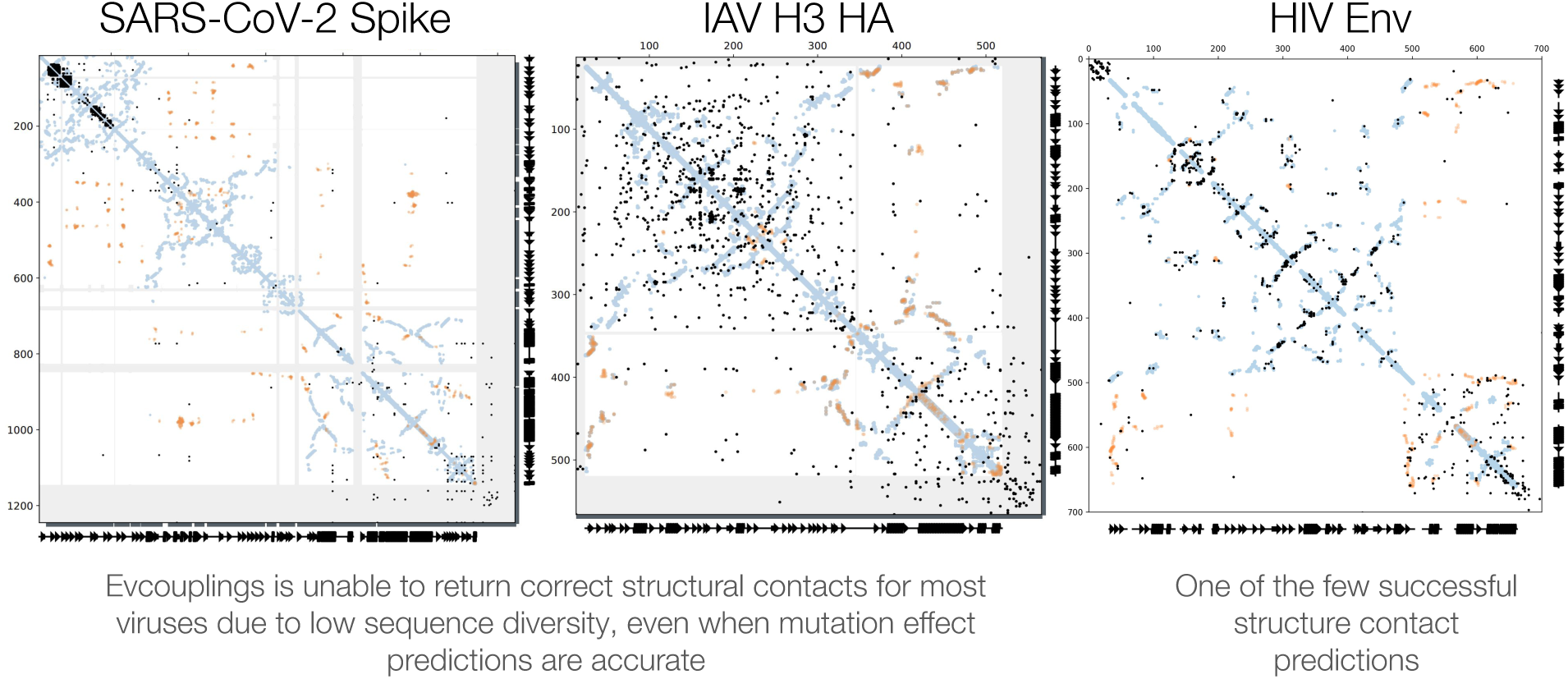
Evolutionary couplings fail to recuperate structure contacts for most viral proteins. The ability to predict structural contacts is a strong predictor of model performance for other taxa. Contact maps from UniRef100 alignments with bit score 0.1 for SARS-CoV-2 Spike, Influenza H3 and HIV Envelope proteins. HIV Env is one of the few examples of viral proteins where evolutionary couplings successfully predict structure contacts. Blue are inter-molecular contacts and orange are intra-molecular contacts from PDB, black are top evolutionary couplings (predicted co-evolving residues pairs) from the Potts model ^135^.

**Supplementary Figure 11:**
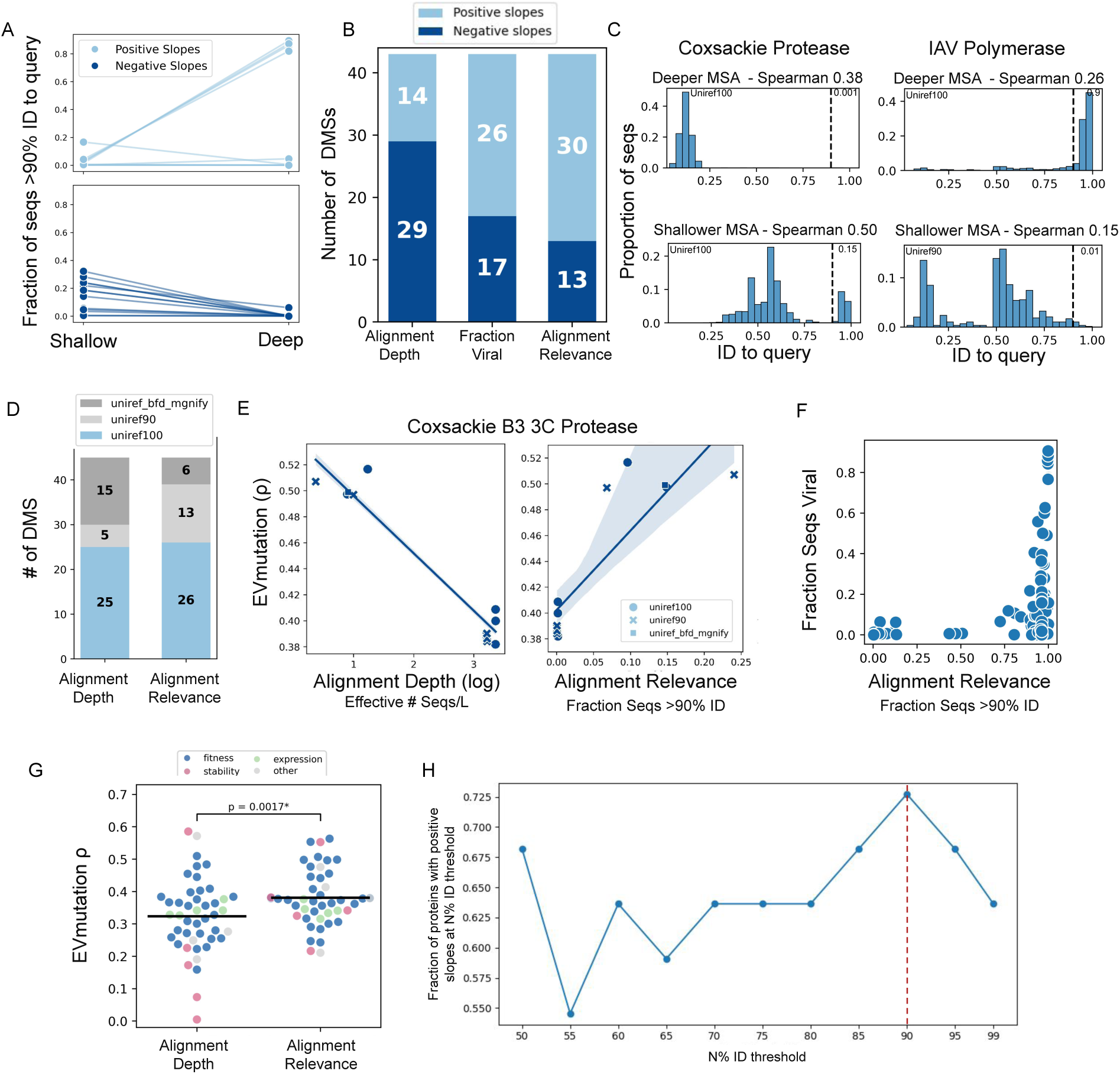
Alignment relevance is a better predictor of model performance than commonly used alignment depth. Alignment relevance is measured as the fraction of sequences in the alignment within greater than 90% identity to the quety. Alignment depth is measured as the number of effective sequences over protein length. (A) Deeper alignments improve (or reduce) performance when a larger (or smaller) proportion of sequences are within 90% identity of the target sequence. Proteins that benefited from deeper alignments had a higher proportion of closely related sequences, while those that performed worse contained more distantly related sequences. (B) Alignment relevance is the best predictor of PSSM performance over alignment depth (Neff / L) and the fraction of viral sequences in the alignment. Reported are the number of DMS assays with positive (light blue) or negative (dark blue) slopes in performance as a function of different alignment metrics. Only proteins with significant differences in performance (Spearman difference between best and worst alignment *>*0.1) are reported. (C) Increasing alignment relevance increases Spearman correlation from 0.38 to 0.50 for the coxsackievirus protease (query). Using a deeper alignment, the majority of sequences have less than 25% identity to query (with less than 0.1% of sequences within 90% identity, dotted line). Using a shallower alignment, 15% of sequences are within 90% identity of the query. Conversely for IAV polymerase, the deeper alignment has a higher relevance and yields higher performance. (D) Fraction of selected alignments using each database for old and new alignment selection metric. (E) Using alignment relevance as a metric for alignment selection instead of depth is a better predictor of Spearman performance for coxsackievirus protease. (F) Relationship between alignment relevance and fraction of viral sequences in the alignment across Uniref100 alignments at different bitscores. (G) Improved model performance (Spearman correlation between Potts model prediction and DMS) across EVEREST DMS benchmark when using alignment relevance versus depth for alignment selection. (H) Model performance at different % ID threshold for defining alignment relevance across a subset of ProteinGym.

**Supplementary Figure 12:**
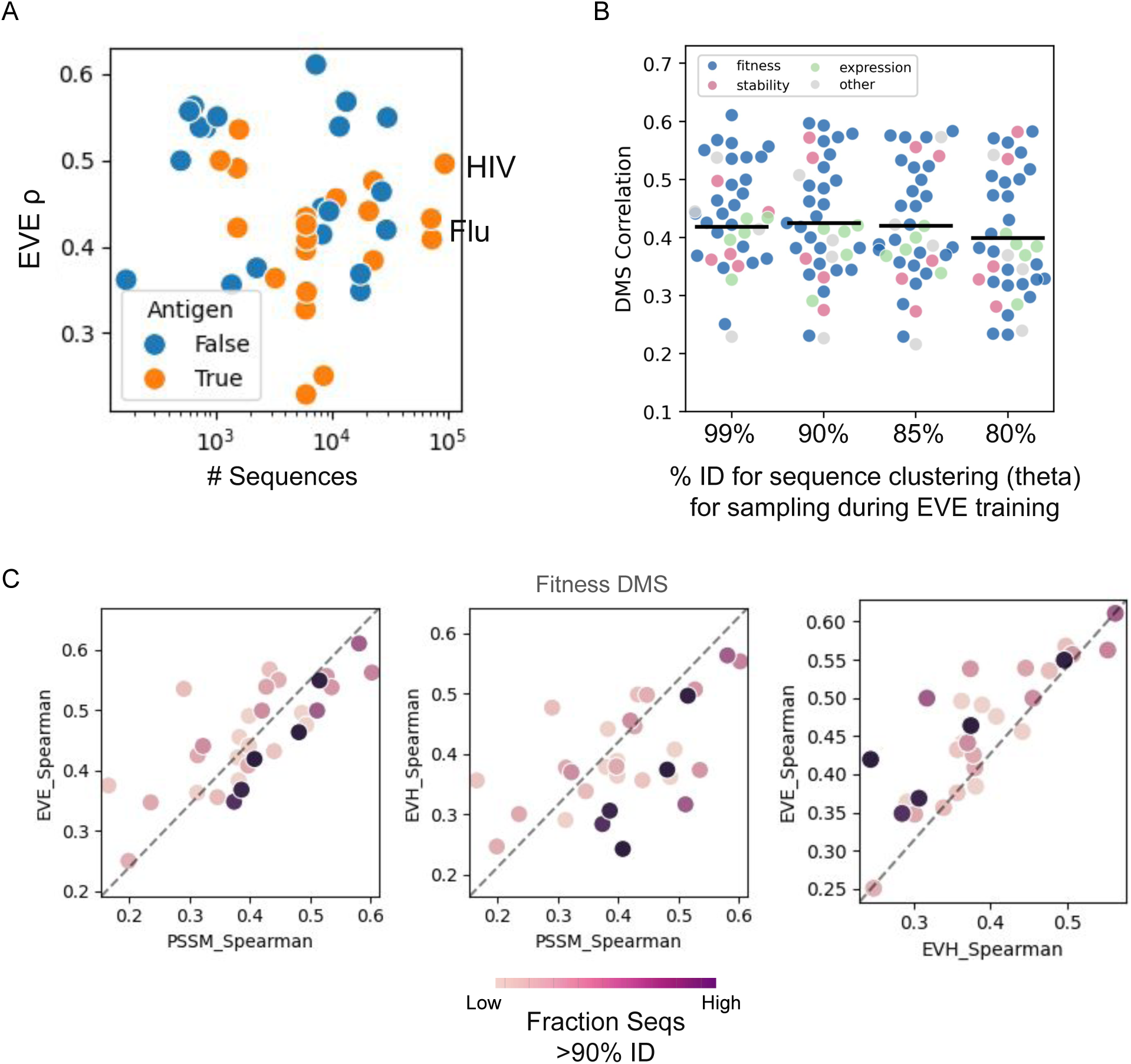
(A) No relationship between raw number of sequences and EVE performance, with no clear performance boost for the most highly sequenced viruses (HIV and Flu) (B) Similar model performance (Spearman correlation between EVE model prediction and DMS) across EVEREST DMS benchmark when using different percentage sequence identity cutoffs (thetas) for clustering to determine sampling frequency during EVE training. (C) Relationship between alignment relevance and performance of different alignment-based models. PSSMs are more likely to outperform Potts models when the alignment is composed of more relevant sequences to the query.

**Supplementary Figure 13:**
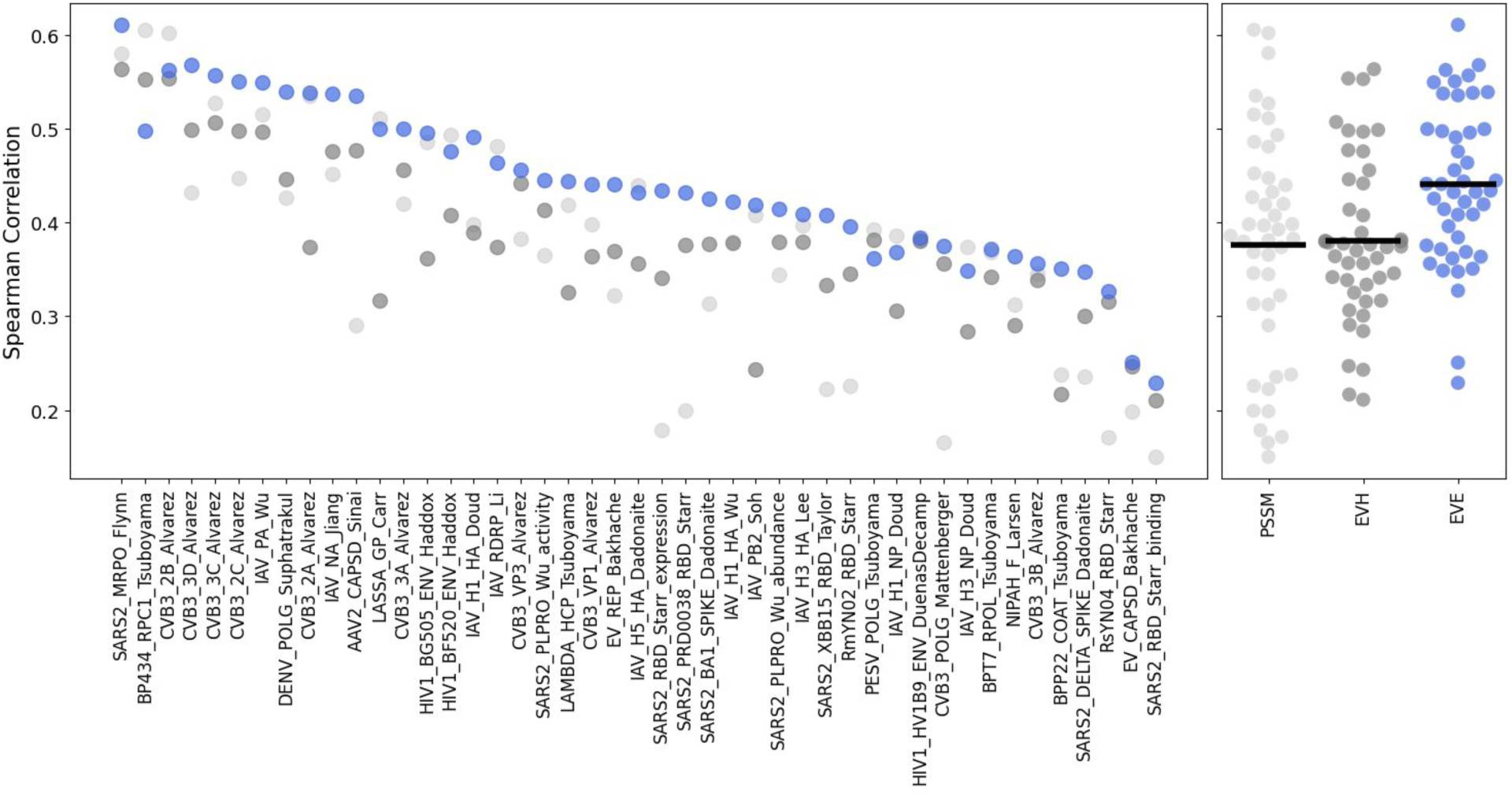
Spearman rank correlation between alignment-based model score and DMS score across EVEREST DMS benchmark. EVE outperforms other alignment-based models.

**Supplementary Figure 14:**
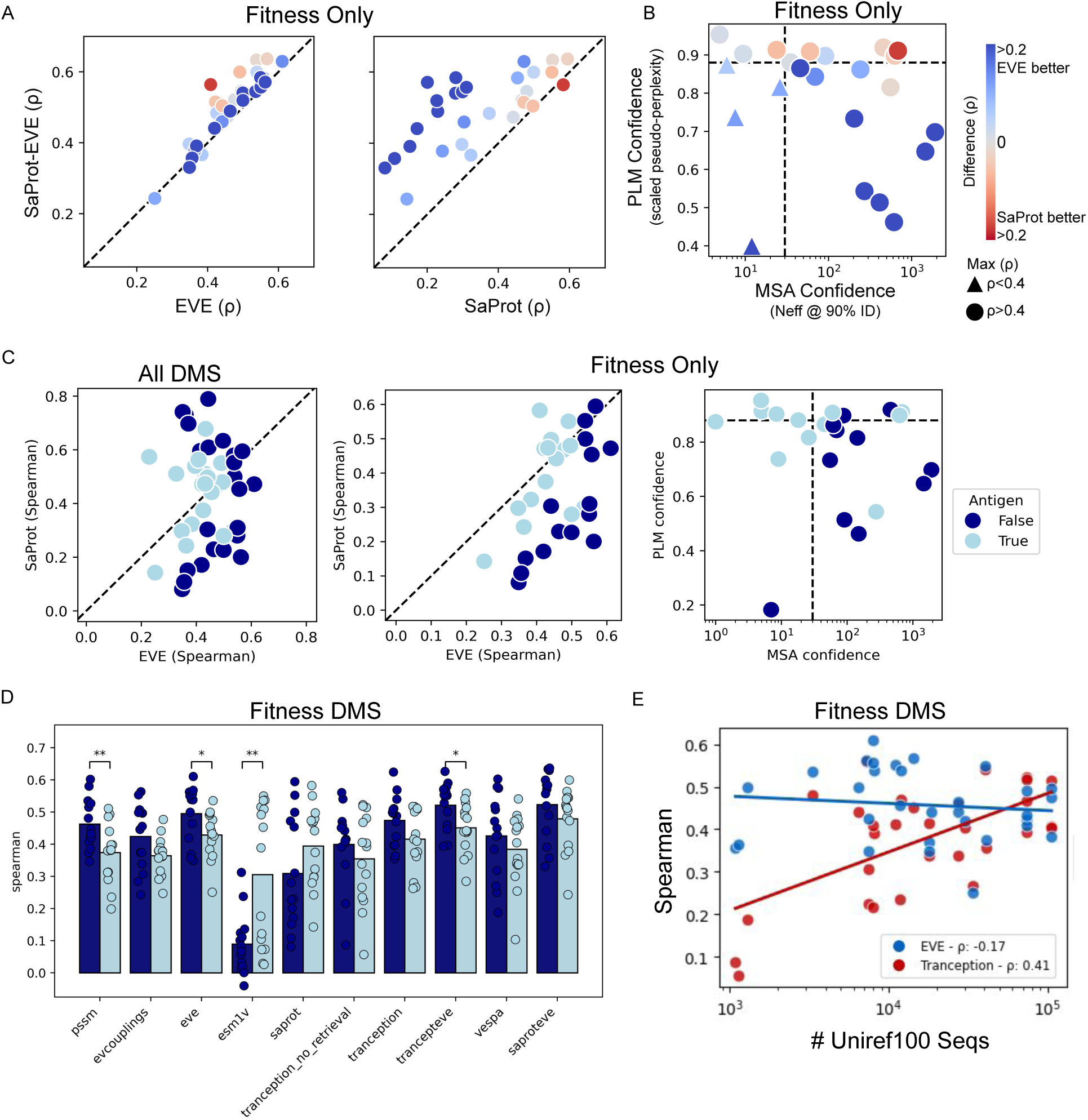
(A) SaProt-EVE performance meets or exceeds EVE or SaProt correlations for all viral fitness DMSs. (B) Viral proteins in fitness DMS with high EVE reliability scores (Neff90) have better performance with EVE; those with higher SaProt reliability scores (scaled pseudo-perplexity) are better modeled with SaProt. Size depends on if max Spearman for DMS across EVE and SaProt is greater than 0.4. Lines shown at optimal thresholds that maximize sensitivity and specificity for whether confidence metric is predictive of model-DMS correlation being higher than 0.4 (high confidence lines). (C) Non-antigens have higher consistently higher EVE scores and MSA confidence values for a particular SaProt score range, whereas the highest SaProt scores are a mix of antigens and non-antigens. (D) Some models have different performances on fitness DMSs for antigens and non-antigens. Significance by paired t-test. (E) Tranception performance increases with more related sequences in Uniref100 by MMseqs search. The associated best EVE model performance shown with respect to the Uniref100 number of sequences, to inform model selection.

**Supplementary Figure 15:**
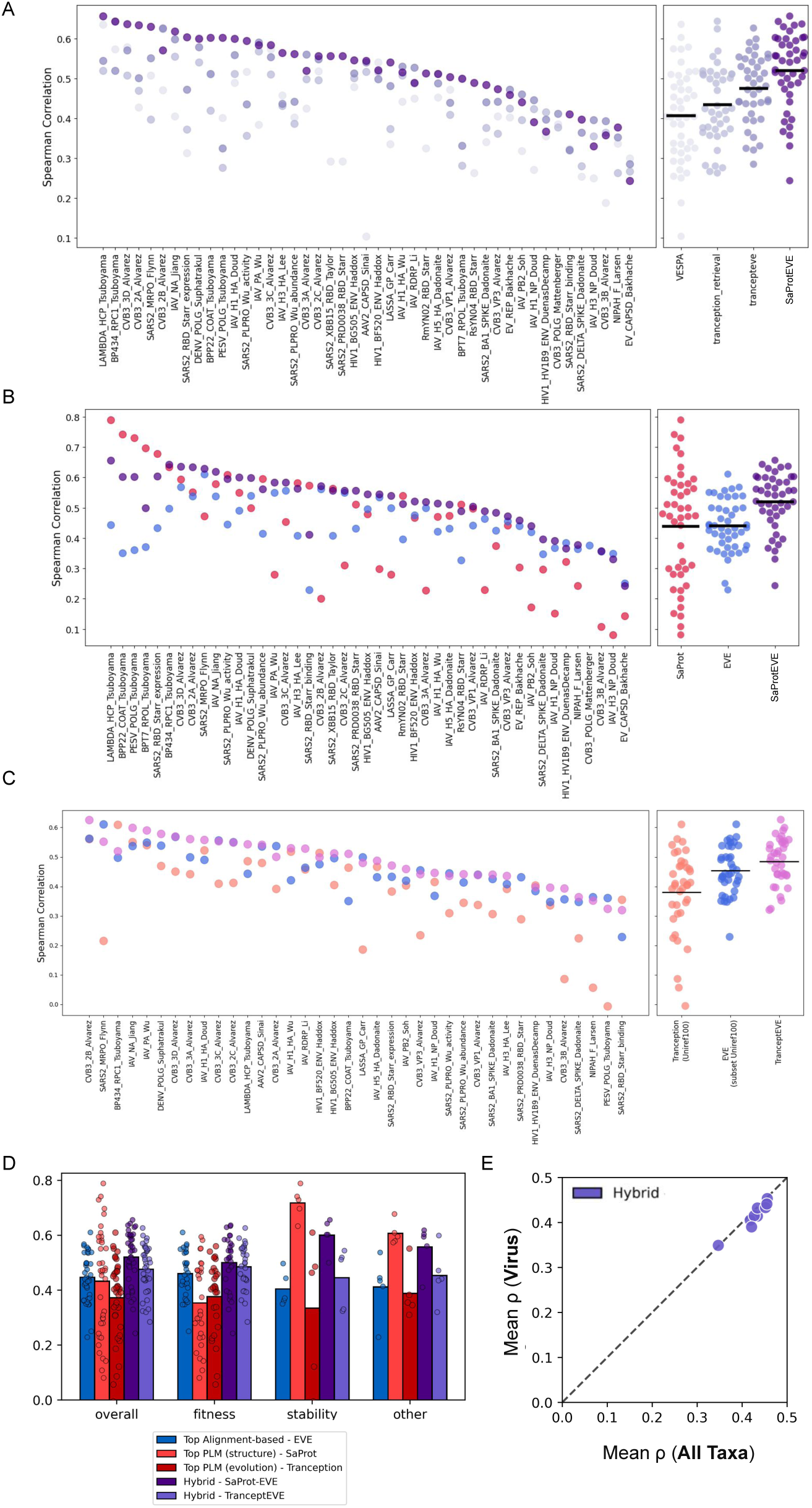
(A-C) Breakdown of hybrid models performance on each protein in EVEREST DMS benchmark. (A) Performance of models shown in main: VESPA, Tranception with MSA, TranceptEVE, and SaProt-EVE. (B) SaProt, EVE and SaProt-EVE (C) Tranception, EVE and TranceptEVE. (D) Alignment-based models, PLMs, and hybrids perform best on different DMS phenotypes. (E) Unlike protein language models, hybrid models have similar performance for viral proteins compared to proteins from other taxa. Performance is measured as Spearman correlation with experimental deep mutational scanning (DMS) assays from prior benchmark ^17^.

**Supplementary Figure 16:**
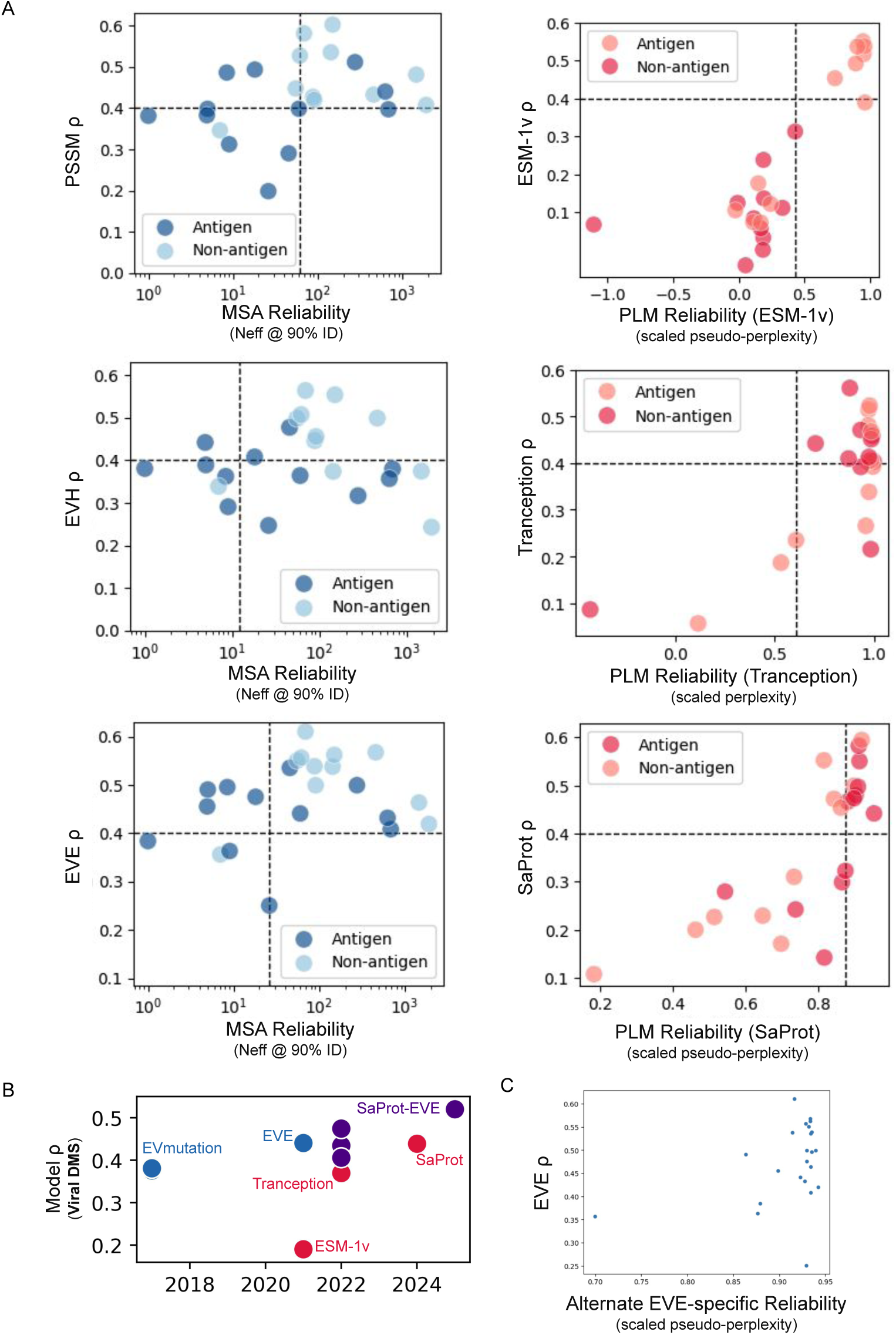
(A) Reliability scores predict performance for a given viral protein for PSSM, EVE, ESM-1v, and SaProt models, though not for Potts model or Tranception. For alignment-based models we use effective number of sequences at 90% sequence identity to the query and for PLMs we use scaled perplexity (autoregressive) or pseudo-perplexity (MLM). The optimal threshold that maximizes sensitivity and specificity for predicting high EVE performance (defined as Spearman *ρ >* 0.4) was 30 (dotted line). For comparability, we subset to the set of fitness assays, with length *<*1024 (context size of PLMs) and date post-2015. See methods for more details. (B) Average spearman on viral-specific DMS benchmarks across evaluated models have improved over time. (C) Correlation between EVE performance and an alternate attempt at a reliability metric using a “pseudo-perplexity” of the EVE Evidence Lower Bound (ELBO), a lower bound to the log-marginal likelihood of each amino acid at each position.

**Supplementary Figure 17:**
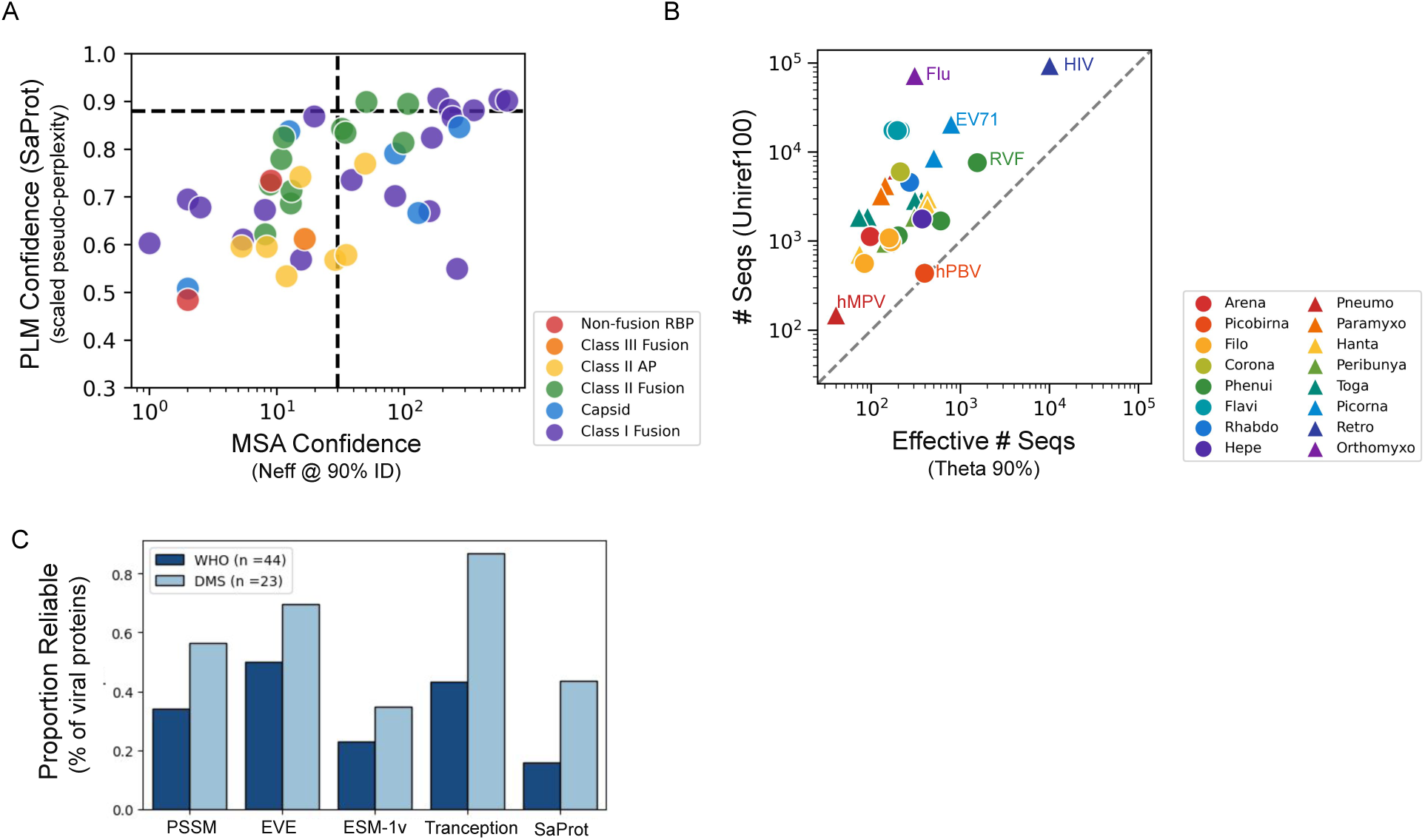
A. PLM (SaProt) and MSA reliability scores for all 44 WHO priority or prototype RNA viral antigens. WHO priority pathogens colored by structural class. B. Number of related sequences in a JackHMMER search of Uniref100 and when clustered with theta 0.9 for each priority virus. HIV has the most sequences, including when looking at diverse effective number sequences at theta 0.9. C. Proportion of reliable viral proteins across viral DMS (subset of 23 comparable fitness assays) or 44 WHO priority or prototype RNA viral antigens. Models included where reliability estimates predict performance (see Fig. S16). DMS benchmarks overestimates generalizability of model performance due to focus on well-characterized viruses.

**Supplementary Figure 18:**
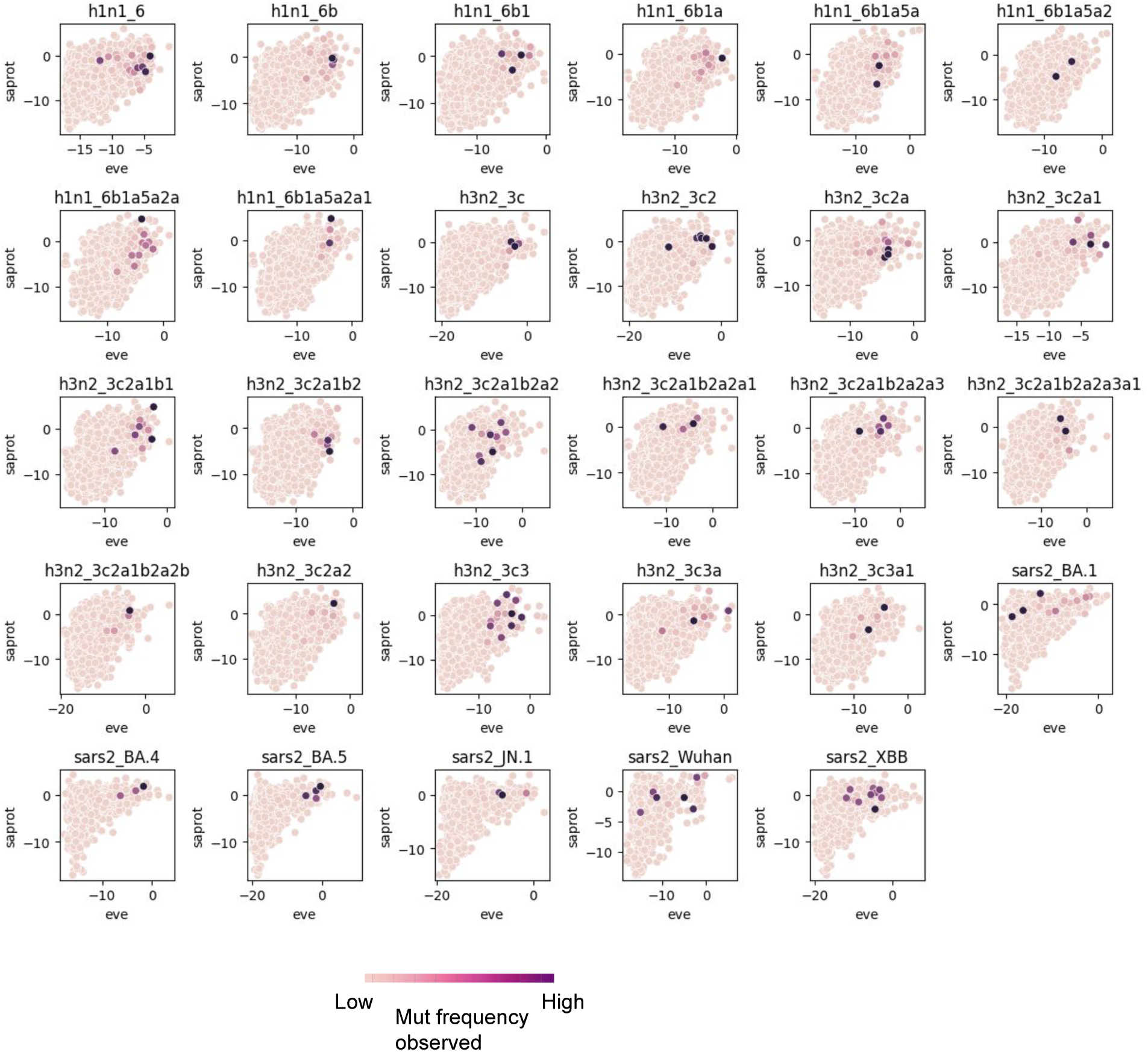
Distribution of model predicted mutation scores for SaProt and EVE colored by mutation frequency.

